# Electron transport chain biogenesis activated by a JNK-insulin-Myc relay primes mitochondrial inheritance in *Drosophila*

**DOI:** 10.1101/642314

**Authors:** Zong-Heng Wang, Yi Liu, Vijender Chaitankar, Mehdi Pirooznia, Hong Xu

## Abstract

Oogenesis features an enormous increase in mitochondrial mass and mtDNA copy number, which are required to furnish mature eggs with adequate mitochondria and to curb the transmission of deleterious mtDNA variants. Quiescent in dividing germ cells, mtDNA replication initiates upon oocyte determination in the *Drosophila* ovary, which necessitates active mitochondrial respiration. However, the underlying mechanism for this dynamic regulation remains unclear. Here, we show that an feedforward insulin-Myc loop promotes mitochondrial respiration and biogenesis by boosting the expression of electron transport chain subunits and factors essential for mtDNA replication and expression, and mitochondrial protein import. We further reveal that transient activation of JNK enhances the expression of insulin receptor and initiates the insulin-Myc signaling loop. Importantly, this signaling relay ensures sufficient mtDNA in eggs and limits the transmission of a deleterious mtDNA mutation. Our study demonstrates cellular mechanisms that couple mitochondrial biogenesis and inheritance with oocyte development.

## INTRODUCTION

Mitochondria host a number of biosynthetic pathways and produce most cellular ATP through oxidative phosphorylation carried out by the electron transport chain complexes (ETC) on mitochondrial inner membrane. While the majority of mitochondrial proteins are encoded on the nuclear genome, synthesized in cytoplasm, and imported to mitochondria, a subset of core ETC components are encoded on the mitochondrial genome (mtDNA) and synthesized inside the mitochondrial matrix. Thus, mitochondria biogenesis and ETC activity in particular, rely on the coordinated expression of both nuclear encoded mitochondrial genes and mtDNA (Falkenberg et al., 2007). To meet the diverse energy demands and metabolic programs of the cell, the abundance of mitochondria and their activity differ in a variety of tissues and developmental processes. Mitochondria are transmitted exclusively through maternal lineage in most metazoans (Wallace, 2008), which demands complex regulations on mitochondrial biogenesis and ETC activity during oogenesis. Animal oocytes are hundreds of times larger than their progenitors (Picton et al., 1998). Accompanying this tremendous cell growth during oogenesis, mitochondria undergo prodigious biogenesis and mtDNA copy number increases over thousands of folds (Stewart et al., 2008). The massive amount of mitochondria in the mature oocyte is necessary to power the early embryonic development, as inadequate mitochondrial contents often lead to embryonic lethality (May-Panloup et al., 2007). However, the mechanism by which the germline couples mitochondrial biogenesis to oocyte development remains elusive.

In addition to furnishing mature oocytes with sufficient amount of mitochondria, oogenesis is also tasked with limiting the transmission of harmful mtDNA mutations. Mitochondrial genome is prone to accumulate mutations due to its close vicinity to the highly-mutagenic free radicals in the mitochondrial matrix and a lack of effective repair mechanisms (Pesole et al., 1999). Yet, harmful mtDNA mutations are rare in populations (Stewart and Larsson, 2014), underscoring the presence of efficient mechanisms to limit their transmission through the female germline. We previously reported that mtDNA replication depends on active respiration in the *Drosophila* ovary (Hill et al., 2014). Healthy mitochondria with wild type genome propagate more vigorously than defective ones afflicted with harmful mutations, which thereby curbs the transmission of deleterious mtDNA mutations into the next generation (Zhang et al., 2019). Therefore, active ETC appears to be a stress test for mtDNA functionality, and is essential for mtDNA selective inheritance. Nonetheless, how the activity of ETC is regulated during oogenesis is not well understood.

Insulin signaling (IIS), an evolutionary conserved pathway that controls cell growth and proliferation (Oldham and Hafen, 2003), has also been shown to regulate ETC biogenesis and ATP production in human skeletal muscles (Stump et al., 2003). In the *Drosophila* ovary, IIS promotes the growth of follicles from early to middle stages of oogenesis (LaFever and Drummond-Barbosa, 2005). Prior to the nurse cell dumping, IIS activity is decreased, which relieves the inhibition on GSK3, and thereby shut down mitochondrial respiration (Sieber et al., 2016). However, oogenesis begins with germline stem cells (GSCs) that are considered not to obtain ATP from respiration (Kai et al., 2005). We predicted there has to be developmental cues to activate mitochondrial respiration in the late germarium stage when mtDNA replication commences. IIS represents a logical candidate to modulate this metabolic transition in early oogenesis. Nonetheless, it remains to be explored how IIS is dynamically regulated during oogenesis and whether it is indeed involved in the aforementioned regulation. Furthermore, little is known regarding how IIS modulates ETC activity and mtDNA biogenesis in general.

In this study, we find that mitochondrial respiration is quiescent in GSCs and dividing cysts, but markedly up-regulated in the late germarium, the same spatial-temporal pattern as mtDNA replication. We uncovered a feedforward loop between IIS and Myc protein which orchestrates the transcriptional activation of respiration and mtDNA replication. Furthermore, transient JNK activity boosts insulin receptor (InR) transcription to enhance the IIS-Myc loop. Our work uncovers how developmental programs couple mitochondrial biogenesis with cell growth and its impact on mitochondria inheritance.

## RESULTS

### Coordinated transcription regulations on both nuclear and mitochondrial genome control ETC biogenesis

mtDNA replication is significantly increased in the post-mitotic germ cells in late germarium and relies on mitochondrial inner membrane potential (Ψ_m_) and ETC activity (Hill et al., 2014). We therefore hypothesized that mitochondrial respiration might be developmentally regulated in a spatial-temporal pattern similar to that of mtDNA replication. To test this idea, we monitored Ψ_m_, which is an indicator of mitochondrial respiration, in the developing germ cells. We found that Ψ_m_, measured as the ratio of TMRM (an indicator for membrane potential) to MitoTracker Green (an indicator for mitochondrial mass) (Zhang et al., 2019), was markedly higher in region 2B than the earlier stages in the germarium (Fig 1A), indicating that respiration is activated in the 16-cell cysts, concomitantly with the onset of mtDNA replication. Consistently, ETC activity, indicated by a dual SDH (succinate dehydrogenase)/COX (cytochrome C oxidase) colorimetric assay (Ross, 2011), was much higher in region 2B than the earlier germarium stages and remained high until stage 10 egg chamber (Fig 1B and S1A). These results suggest that ETC activity is up-regulated in the late germarium stages.

**Figure 1.**
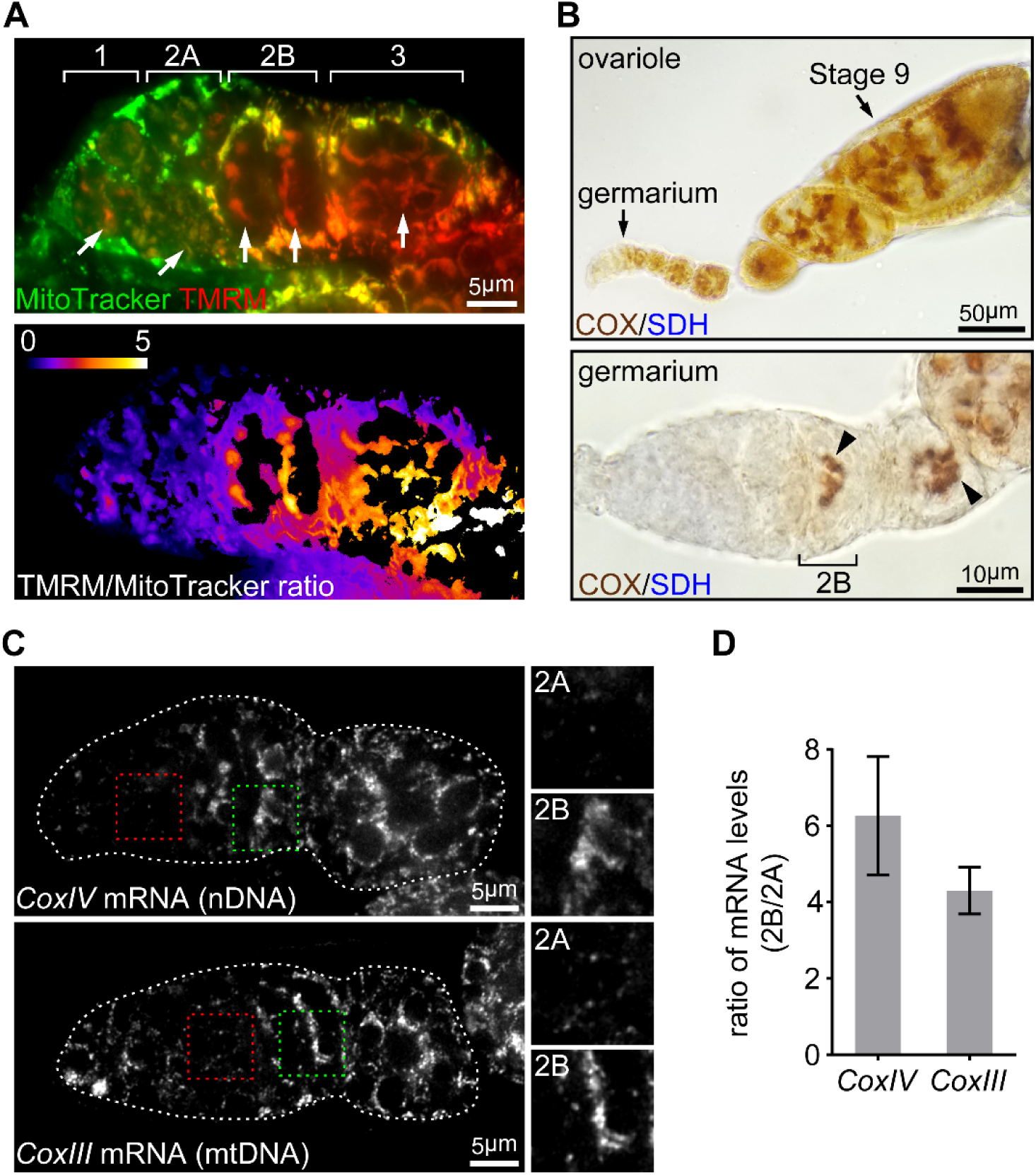
ETC activity and gene expression sharply increase at germarium stage 2B. (A) Upper panel: a representative image of a germarium stained with TMRM (a membrane potential marker) and MitoTracker Green (a mitochondrial mass marker). Germarium regions are indicated. Arrows indicate mitochondria in germ cells (GCs). Lower panel: TMRM/MitoTracker Green ratiometric image, indicating that mitochondria membrane in in stem cells and dividing cysts is low, but markedly increased in 16 cell cysts and budding egg chambers. (B) Upper panel: a representative image of a wt ovariole (from germarium to stage 9 egg chamber) stained for the COX/SDH dual activities. Lower panel: a representative high-magnification image of a germarium stained for COX/SDH. Note the onset of COX/SDH activity in region 2B of the germarium (arrowhead). (C) Visualization of the *CoxIV* and *CoxIII* mRNA in germaria from wt flies with fluorescently labelled DNA probes by FISH. Germaria are outlined with dotted lines. Right panels illustrate the enlarged areas of germarium region 2A (red dotted line) and 2B (green dotted line), respectively, shown in left panels. (D) Quantification for the relative expression level of *CoxIV* or *CoxIII* mRNA in different regions of germarium. Note that both transcripts are markedly induced in region 2B germarium.

We next asked whether the dynamic pattern of ETC activity in the germarium reflected the expression of ETC subunits. Except for complex II (SDH) that are encoded on nuclear genome only, all other ETC genes are encoded by both nuclear and mitochondrial genomes. Thus, we performed fluorescence *in situ* hybridization (FISH) with RNA probes specific to mRNAs of either nuDNA- or mtDNA-encoded ETC subunits in ovaries. Both COXIV (nuclear-encoded) and COXIII (mtDNA-encoded) transcripts exhibited low expression in earlier regions, but increased 4 to 6 folds in region 2B, recapitulating the pattern of ETC activity (Fig 1C and 1D). The same pattern was observed for Cyt-C1 (nuclear-encoded) and Cyt-B (mtDNA-encoded) (Fig. S1B). These results confirm that the increased ETC activity detected with our COX/SDH colorimetric assay indeed reflects an increase of ETC genes expression at region 2B germarium. Taken together, these data suggest that the activation of respiration at stage 2B relies on the coordinated transcription of nuDNA- or mtDNA-encoded genes.

### A candidate RNAi screen for upstream regulators of ETC biogenesis

To uncover the upstream developmental cues that initiate ETC gene transcription in the late germarium, we screened a collection of 132 RNAi lines directed at major developmental signaling and factors involved in cellular metabolism and mitochondrial functions (Basson, 2012; Claveria and Torres, 2016; Desvergne et al., 2006; Perrimon et al., 2012). We expressed dsRNAs in the germ cells using a *nanos*-Gal4 (*nos*-Gla4) driver and applied the COX/SDH assay as an indirect measure of ETC abundance (Table S1). We also included a few RNAi lines directed at COX components or genes essential for mitochondrial biogenesis as positive controls. As expected, knock down of these genes consistently impaired ETC activity (Fig 2A). Overall, 6 RNAi lines from the list caused germline degeneration and 12 lines led to reduced ETC activity without causing the loss of germline or defects in development (Fig S2A). Among these 12 lines are components of the IIS/TORC1 signaling, the JNK pathway, cell adhesion molecules, translation regulators and one transcription factor. Notably, all hits impaired activities of both COX and SDH, except for *coxV* RNAi that disrupted COX activity only, indicating that the recovered genes are required for the expression of both nuclear and mitochondrial genes.

**Figure 2.**
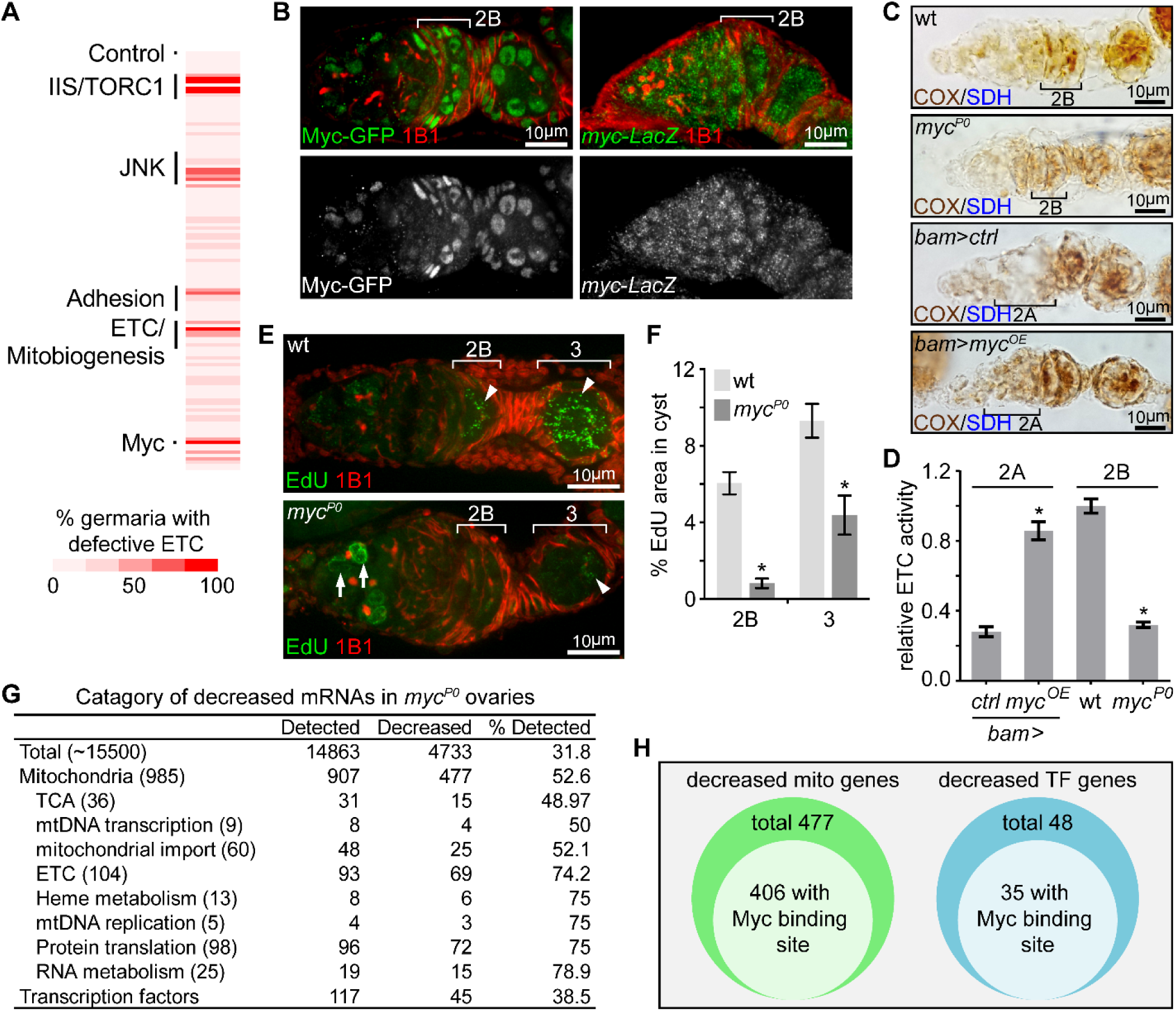
A candidate RNAi screen reveals Myc as an essential regulator for mitochondrial biogenesis. (A) RNAi screening for genes that are required for inducing ETC activity in germaria. For each line, the impact on ETC activity is scored as the percentage of germaria with reduced COX/SDH staining. (B) Left panel: a germarium from flies endogenously expressing Myc-GFP stained with anti-GFP (Green) and anti-1B1 (Red). Right panel: a germarium flies expressing LacZ driven by *myc* endogenous promoter stained with anti-β-galactosidase (Green) and anti-1B1 (Red). Myc protein is expressed in low level in GSCs and dividing cysts, but markedly induced from region 2B germ cells. In contrast, Myc promoter activity is uniform in the germarium. (C) ETC activity in germaria from wt, *myc*^*P0*^, *bam*>*ctrl*, and *bam*>*myc*^*OE*^ ovaries visualized by COX/SDH staining. ETC activity is significantly reduced in the *myc*^*P0*^ mutant, but is ectopically induced when Myc is over-expressed in region 2A by *bam*-Gal4. (D) Quantification of relative ETC activity in germarium regions from ovaries with indicated genotypes. ETC activity is normalized to that in the wt 2B cysts. (E) Visualization of mtDNA replication in germaria from wt and *myc*^*P0*^ ovaries with EdU incorporation (Green) and co-stained with anti-1B1 (Red). Arrows indicate mtDNA replication, while arrowheads point out EdU incorporation into the nuclear genome. (F) Area of EdU puncta (pixels) is normalized to total pixels germline cysts at indicated germarium stages from wt and *myc*^*P0*^ ovaries. (G)Table of genes functioning in mitochondrial processes with at least 3-fold decreased expression in *myc*^*P0*^ ovaries compared with wt ovaries. (H) Diagrams of decreased genes encoding mitochondrial processes and TFs in the *myc*^*P0*^ ovaries. A number of genes in either category has Myc binding site in their regulatory region. Error bars represent SEM. *p < 0.005.

### Myc controls ETC biogenesis and mtDNA replication

The transcription factor Myc emerged as one of the strongest hits from our screen. Additionally, we found that Myc’s expression pattern, monitored with a Myc-GFP fusion protein (Greer et al., 2013), mirrored the pattern of ETC activity in the ovary: low in the early stages, but elevated in germarium region 2B and remaining high until mid-stage egg chambers (Fig 2B and S2B). These observations spurred us to explore the potential roles of Myc in the activation of ETC genes expression at stage 2B. To confirm the result of the Myc RNAi experiment, we utilized a hypomorphic Myc allele, *myc*^*P0*^, which has reduced level of *myc* mRNA, but progresses through early oogenesis (Johnston et al., 1999; Quinn et al., 2004). Consistent with the RNAi result, the *myc*^*P0*^ mutation led to a reduction in ETC activity and mtDNA replication (Fig 2C-2F). Next, to test whether Myc is sufficient for ETC activity induction, we utilized *bam*-*Gal4* to over-express myc ORF in the dividing cyst in region 2A that normally have low levels of both ETC activity and Myc protein. Over-expression of *myc* in this region ectopically enhanced ETC activity (Fig 2C and 2D). Thus, Myc is both necessary and sufficient to stimulate mitochondrial respiration in the ovary.

To gain insight into how Myc regulates mitochondrial biogenesis, we compared the transcriptomes of wt and *myc*^*P0*^ mutant ovaries (Table S2). RNA sequencing (RNAseq) showed that nearly one third of detected transcripts were reduced in *myc*^*P0*^ mutant compared to wt (fold change>3.0, FDR<0.05%) (Fig 2G), consistent with the notion of Myc as a general transcription activator (Orian et al., 2003). Importantly, nuclear-encoded mitochondrial genes were enriched among the down-regulated genes. About 52% of the total mitochondrial genes, and 75% of ETC genes and factors for mtDNA replication and expression were decreased. Myc directly regulates the expression of its targets by binding to a short sequence, CACGTG (E-box) in the regulatory region. Interestingly, 406 out of 477 down-regulated mitochondrial genes have predicted Myc binding sites in their regulatory regions, further substantiating a role of Myc in promoting mitochondrial biogenesis *via* boosting the transcription of mitochondrial genes (Fig 2H). Additionally, 48 transcriptional factors, 35 of which have E-boxes in their regulatory regions, were also decreased in *myc*^*P0*^ mutant ovaries, suggesting that secondary transcriptional controls might also be involved in Myc’s regulation on mitochondrial biogenesis.

### IIS regulates Myc post-transcriptionally through Sgg and Thor

Having identified Myc as the master regulator of ETC biogenies and mtDNA replication in the ovary, we sought to explore how the spatial-temporal pattern of Myc protein was established. Myc can be regulated either transcriptionally or post-transcriptionally by a variety of upstream signals (Gallant, 2013). We first examined *myc* transcription by visualizing its promoter activity using a *myc*-*LacZ* transgene (Neto-Silva et al., 2010). In contrast to Myc protein, which was markedly up-regulated at region 2B, *myc* transcript appeared to be uniform in the germarium (Fig 2B), suggesting that post-transcriptional regulations are responsible for the spatial pattern of Myc protein. IIS/TORC1 signaling is known to regulate both translation and protein stability (Garofalo, 2002; Maurer et al., 2014; Pan et al., 2004), and multiple genes in the IIS/TORC1 signaling emerged from the initial RNAi screen. Consistently, ETC activity and mtDNA replication was severely impaired in ovaries of *chico* mutant flies (Fig 3A-3C), which obtained by combing two *chico* mutant alleles, *chico*^*1*^ (loss of function) and *chico*^*KG*^ (hypomorphic) (Bohni et al., 1999; Song et al., 2010). These data support a critical role of IIS in ETC biogenesis and mtDNA replication. Intriguingly, the activity of IIS, revealed by staining for phosphorylated AKT (p-AKT) (Parker and Struhl, 2015), was also increased in the germarium region 2B and maintained until mid-stage egg chambers (Fig 3D and S3A), a pattern similar to that of Myc protein. In contrast, total AKT staining was uniform in the germarium (Fig 3D). These observations suggest that Myc may be regulated by IIS. Indeed, Myc protein was strongly reduced in *chico*^*1*/*KG*^ mutant ovaries (Fig 3E and 3F). In *chico* RNAi ovaries, Myc protein was also diminished in germ cells, while the expression of *myc*-*LacZ* was not affected (Fig 4B and S4A). Importantly, over-expressing Myc in the *chico* RNAi background largely restored ETC biogenesis in the ovary (Fig 4D). Altogether, these results suggest that up-regulation of IIS in late germarium stimulates ETC biogenesis and mtDNA replication through post-transcriptional control of Myc level.

**Figure 3.**
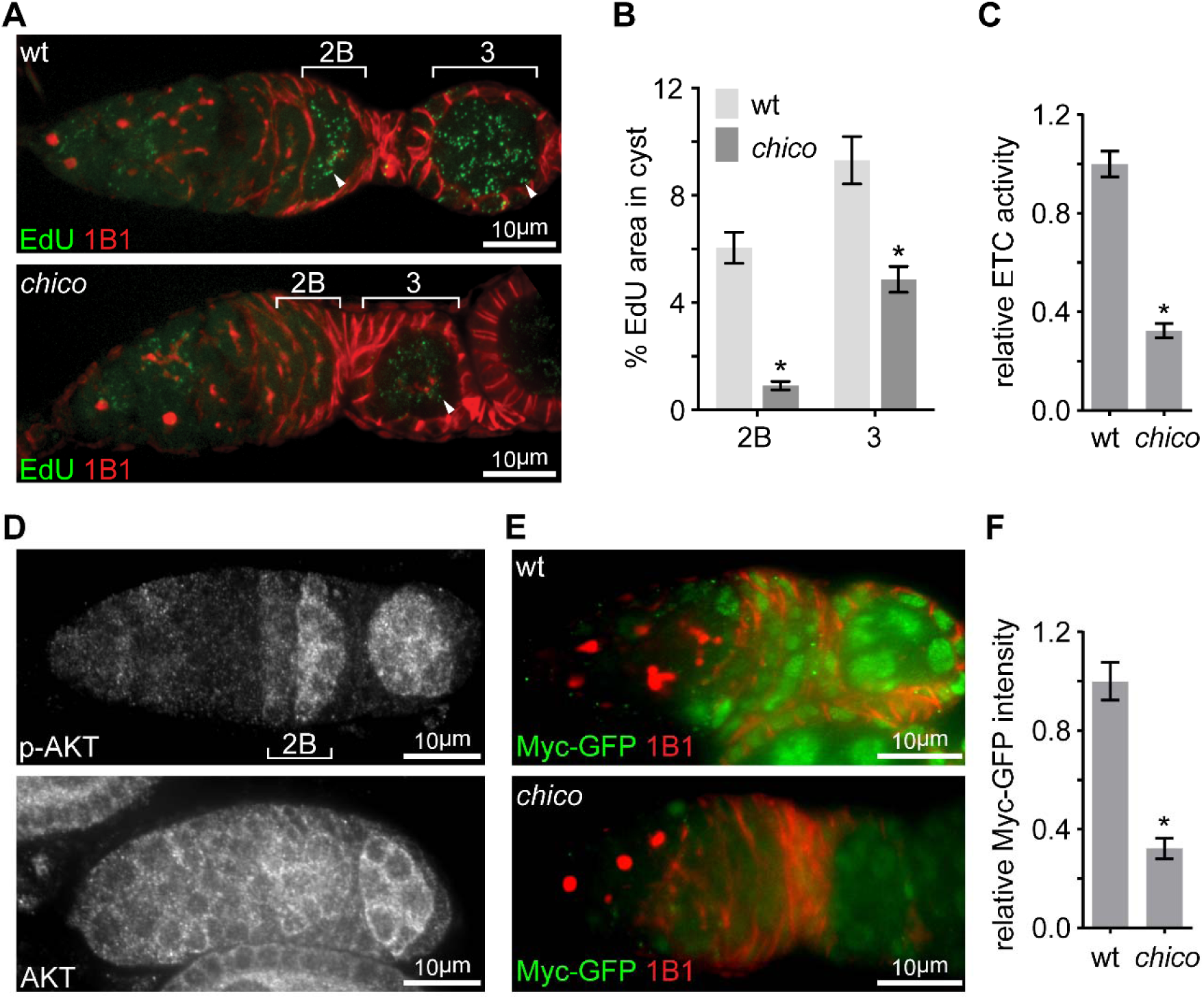
Elevated IIS in late germarium induces Myc protein to prime mtDNA replication and mitochondrial respiration. (A) Representative germaria from wt or *chico*^*1*/*KG*^ mutant flies incorporating EdU stained with anti-1B1. Arrowheads indicate EdU incorporation into mtDNA of germ cells. (B) Quantification of the percentage of EdU area in germline cysts from wt or *chico*^*1*/*KG*^mutant flies. (C) Quantification of relative ETC activity in region 2B cysts from wt or *chico*^*1*/*KG*^ mutant flies. (D) Germaria from wt ovaries stained with anti-AKT and anti-p-AKT, respectively. p-AKT staining is low in both GSCs and dividing cysts, but increased from region 2B germ cells. In contrast, AKT staining is uniform in the germarium. (E) Germaria from wt or *chico*^*1/KG*^ mutant ovaries endogenously expressing Myc-GFP stained with anti-GFP, anti-1B1, and DAPI. Germarium region 2B is underlined with dotted lines. (F) Quantification of relative Myc-GFP intensity in germarium region 2B from wt or *chico*^*1/KG*^ mutant ovaries. Error bars represent SEM. *p < 0.005.

**Figure 4.**
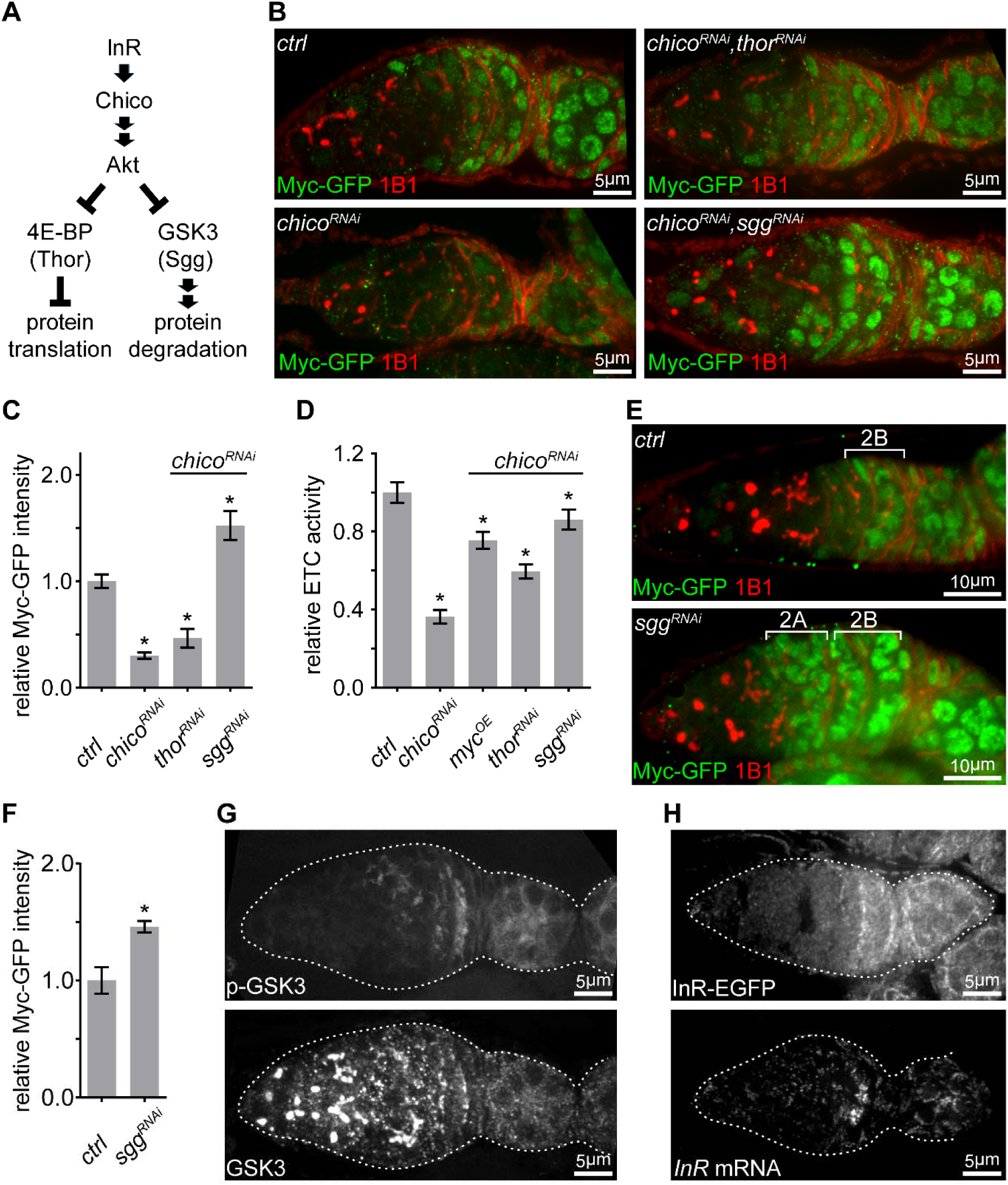
IIS promotes ETC activity via inhibition of GSK3 to stabilize Myc. (A) Schematic of the conserved IIS that inhibits 4E-BP/Thor and GSK3/Sgg to promote protein translation and suppress protein degradation, respectively. (B) Germaria from *nos*>*ctrl*; *nos*>*chico*^*RNAi*^; *nos*>*chico*^*RNAi*^, *thor*^*RNAi*^; and *nos*>*chico*^*RNAi*^, *sgg*^*RNAi*^ ovaries endogenously expressing Myc-GFP stained with anti-GFP (green) and anti-1B1 (red). Germ cells from region 2B are outlined with dotted lines. (C, D) Quantification of Myc-GFP intensity (C) and ETC activity (D) in germarium region 2B from ovaries with indicated genotypes, normalized to the intensity or activity values in germaria with *ctrl* over-expression. (E) Germaria from *nos*>*ctrl* and *nos*>*sgg*^*RNAi*^ ovaries endogenously expressing Myc-GFP stained with anti-GFP and anti-1B1. Myc protein is up-regulated in both region 2A and region 2B germ cells in the *sgg* RNAi ovary. (F) Quantification of relative Myc-GFP intensity in germarium region 2B from *nos*>*ctrl* and *nos*>*sgg*^*RNAi*^ ovaries. Myc-GFP intensity is normalized to that of region 2B cysts with *ctrl* over-expression. (G) A germarium from wt flies stained with anti-GSK3 and anti-p-GSK3. Germaria are outlined with dotted lines. (H) Upper panel: a representative image of a germarium from ovaries expressing endogenous InR-EGFP. Lower panel: visualization of the *InR* mRNA in germarium with fluorescently labelled DNA probes by FISH. Germaria are outlined with dotted lines. Error bars represent SEM. *p < 0.005.

Next, we explored how IIS regulates Myc. IIS either promotes protein translation by repressing 4E-BP/Thor, or stabilizes its targets by antagonizing GSK3/Sgg dependent protein degradation (Fig 4A) (Garofalo, 2002; Maurer et al., 2014; Pan et al., 2004). Knocking down of *sgg* in a *chico* RNAi background restored Myc protein level and ETC activity, while *thor* RNAi only partially rescued both (Fig 4B-4D). Intriguingly, *sgg* RNAi not only elevated Myc protein level in region 2B and thereafter, but also strongly induced Myc in earlier stages where Myc protein is not normally present (Fig 4E and 4F). Next, we examined the pattern of GSK3 and GSK3 activity in the germarium. GSK3 activity is suppressed by IIS through AKT phosphorylation on GSK3 Ser9. GSK3 protein visualized by both antibody staining and an endogenous expressed Sgg-GFP was ubiquitous throughout oogenesis (Fig 4G and S4F). However, phosphorylated GSK3, the inactive form of GSK3, became evident in region 2B germarium and later stages egg chambers, the same pattern as p-AKT, Myc, and ETC biogenesis (Fig 1B, 3F, 4G, and S4C). Taken together, these data suggest that Sgg is the main regulator of Myc and acts downstream of IIS.

### InR expression is boosted at region 2B germarium

So far, our data has established Myc as the link between IIS, a major pathway regulating cell proliferation and growth, and mitochondrial biogenesis in ovaries. The IIS regulates germ cells growth and proliferation in response to insulin-like peptides (dilps) produced by neuroendocrine cells (LaFever and Drummond-Barbosa, 2005). *Drosophila* has an open circulatory system. In a given tissue, all cells are exposed to a similar level of dilps circulating in the hemolymph. However, instead of being uniform in the germarium, the activity of IIS, indicated by both p-AKT and inhibitory phosphorylated Sgg staining (Fig 3D and 4G), demonstrated a distinct spatial-temporal pattern similar to that of Myc, ETC expression and mtDNA replication. Therefore, some IIS components downstream of dilps are likely differentially expressed in germarium.

To test this idea, we generated an InR-EGFP reporter line by inserting an EGFP at the C-terminus of the *InR* genomic locus. Using this line and other reporter lines (Nagarkar-Jaiswal et al., 2015; Orme et al., 2006; Sarov et al., 2016), we examined the expression patterns of InR and other components of IIS signaling upstream of AKT. InR-EGFP was upregulated in region 2B (Fig 4H), while all other components in the IIS signaling examined were ubiquitously expressed in the germarium (Fig S5A). Additionally, *InR* mRNA, visualized by FISH, demonstrated the same pattern as that of InR-EGFP (Fig 4H), suggesting that up-regulation of *InR* transcription enhances IIS to boost mitochondrial biogenesis.

### Both JNK pathway and Myc promote IIS activity via *InR* transcription

We next asked how *InR* transcription was elevated at region 2B germarium. The JNK pathway, which transcriptionally controls various cellular processes (Weston and Davis, 2007), had emerged from the initial RNAi screen. Consistent with the RNAi screen, homozygous *bsk*^*1*^ clones showed impaired mtDNA replication, compared to *bsk*^*1*^/*+* heterozygous germ cells (Fig 5A and 5B). Interestingly, JNK signaling activity, visualized by a *puc*-*LacZ* reporter (Martin-Blanco et al., 1998), was sharply up-regulated in late germarium stages, but deceased and eventually disappeared in growing egg chambers (Fig 5C and S5B). The partial overlap between the spatial patterns of IIS and JNK activation, and the phenotypic resemblance between IIS and JNK mutations on ETC activities and mtDNA replication, suggest a potential link between these two pathways. Indeed, IIS activity, Myc protein, and *InR* mRNA were all markedly reduced in ovaries expressing dsRNA against either *bsk* or *jra*, fly homolog of JNK or Jun, respectively (Fig 5D-5I and S5C-S5E). In *bsk* RNAi or *jra* RNAi background, over-expression of *InR* restored Myc level and over-expression of either *InR* or *Myc* rescues ETC activity (Fig 5H and 5I). In contrast, enhancing JNK signaling by *puc* RNAi failed to rescue defective ETC activity in a *chico* RNAi background (Fig 5I). Together, these observations suggest that JNK promotes ETC biogenesis in late germarium stages thorough boosting IIS.

**Figure 5.**
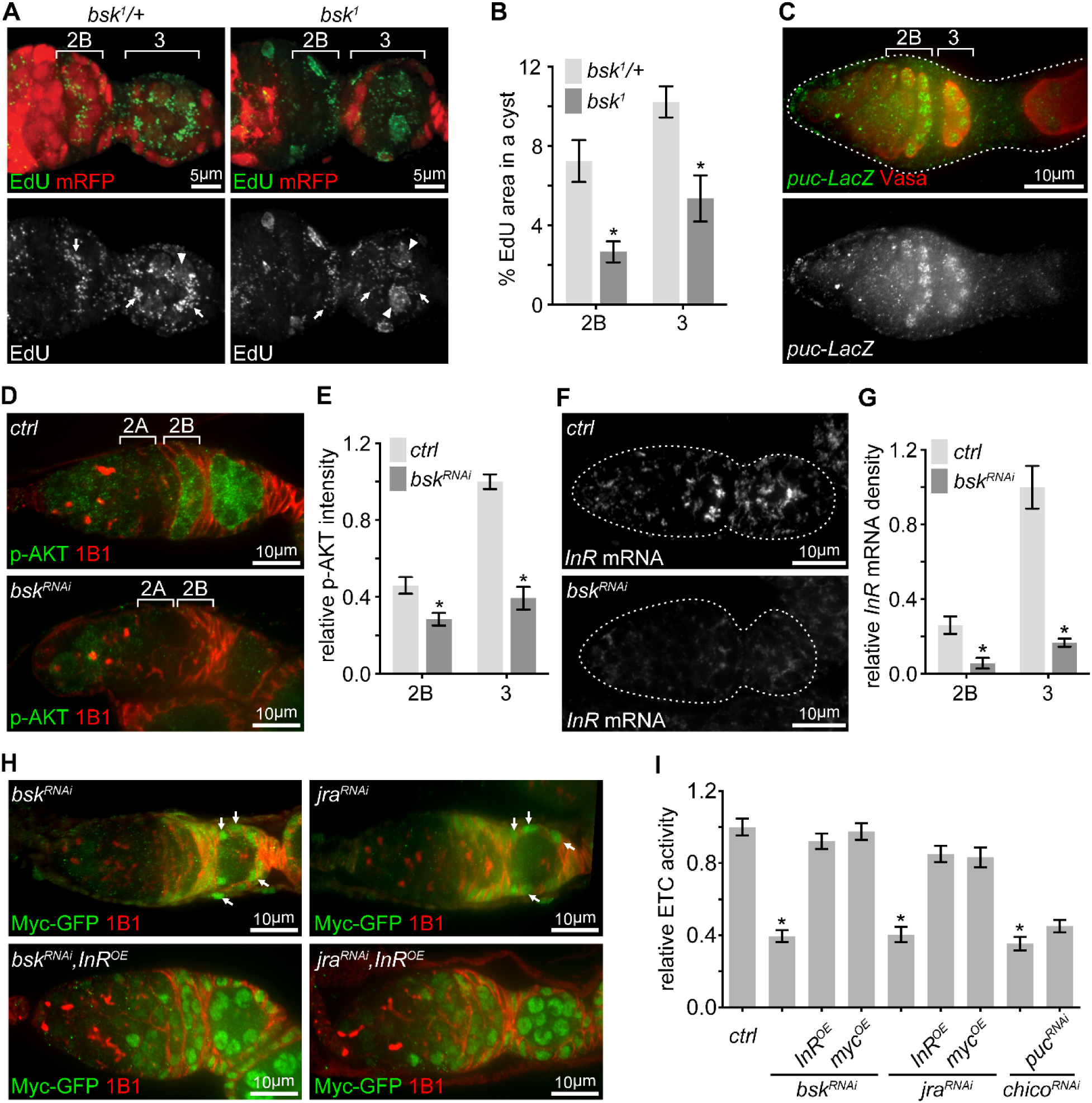
Transient JNK activation in the germarium boosts the IIS-Myc signaling. (A) Representative germaria with *bsk*^*1*^ FRT clones with EdU incorporation (green) to visualize mtDNA replication. *bsk*^*1*^*/+* cells are positive for mRFP (red), while *bsk*^*1*^ mutant cells are negative for mRFP. Arrows indicate EdU incorporated into mtDNA, while arrowheads point out EdU incorporated into the nuclear genome. (B) The percentage area of EdU incorporated into mtDNA in germline cysts at indicated germarium stages from *bsk*^*1*^*/+* and *bsk*^*1*^ clones. (C) Germarium from ovaries expressing LacZ driven by the *puc* promoter stained with anti-β-galactosidase (green) and anti-Vasa (red). Germarium is outlined with dotted lines. (D) Germaria from *nos*>*ctrl* and *nos*>*bsk*^*RNAi*^ flies stained with anti-p-AKT (green) and anti-1B1 (red). Note that IIS activity is markedly reduced when JNK signaling is decreased by *bsk* RNAi. (E) Quantification of p-AKT intensity from cysts in germarium region 2A and 2B of ovaries with indicated genotypes. p-AKT intensity is normalized to that of region 2B cysts from the *ctrl* line. (F) Visualization of the *InR* mRNA by FISH in germaria from *nos*>*ctrl* and *nos*>*bsk*^*RNAi*^ ovaries co-stained with DAPI. Note that *InR* mRNA level is decreased by *bsk* RNAi. Germaria are outlined with dotted lines. (G) Quantification of *InR* mRNA density in region 2B germ cells from ovaries with indicated genotypes. (H) Germaria from *nos*>*bsk*^*RNAi*^; *nos*>*bsk*^*RNAi*^, *InR*^*OE*^; *nos*>*jra*^*RNAi*^; *nos*>*jra*^*RNAi*^, *InR*^*OE*^ flies endogenously expressing Myc-GFP co-stained with anti-GFP and anti-1B1. Note that Myc-GFP level in germ cells is markedly lower than that in follicle cells pointed out by arrows. (I) ETC activity in region 2B cysts from ovaries with indicated genotypes. ETC activity is normalized to that of region 2B cysts from the *ctrl* line. Error bars represent SEM. *p < 0.005.

IIS was elevated in region 2B and remained active until stage-10 egg chambers, the same period during which ETC biogenesis and mtDNA replication are active. However, JNK is only transiently activated at region 2B. Therefore, additional regulations must be involved to maintain IIS activity after JNK activity subsides. Our RNAseq results showed that *InR* mRNA was down-regulated in the *myc*^*P0*^ ovary compared with controls (Table S2), suggesting that Myc might activate *InR* transcription. Indeed, both *InR* mRNA level measured by FISH and IIS activity indicated by p-AKT were reduced in *myc*^*P0*^ mutant or *chico* RNAi ovaries (Fig 6A-6D, S6A, and S6B). Significantly, over-expression of *myc* using *bam*-Gal4 in region 2A ectopically induced *InR* transcription and IIS activity (Fig 6A-6D, S6A, and S6B), suggesting that Myc can indeed enhance IIS by boosting *InR* expression. Together, our results highlight an IIS-Myc positive feedback loop that promotes ETC biogenesis and mtDNA replication in the ovary.

**Figure 6.**
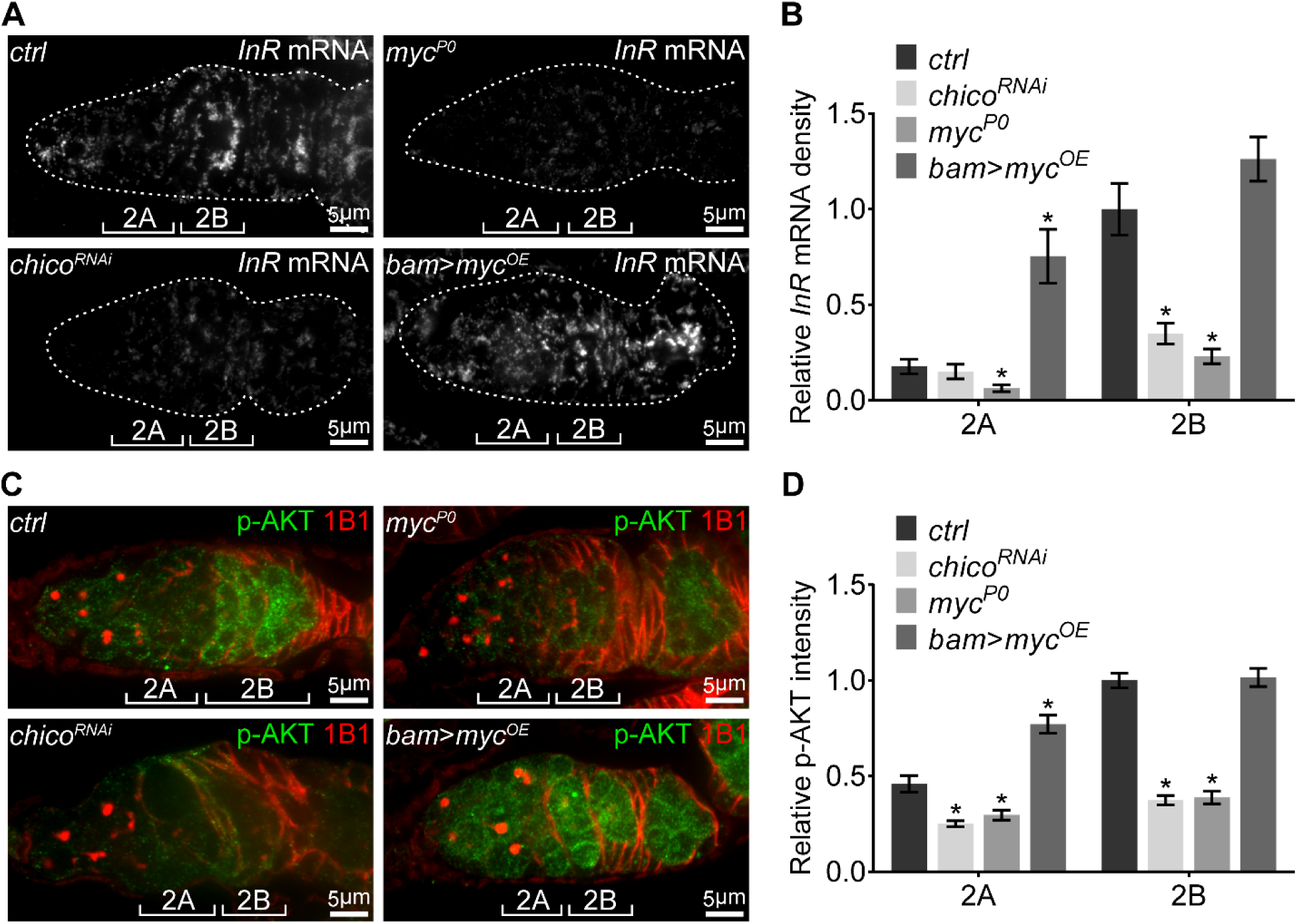
A positive feedback regulatory loop between IIS and Myc. (A) Visualization of the *InR* mRNA with fluorescently labelled DNA probes (green) by FISH in germaria from *nos*>*ctrl, nos*>*chico*^*RNAi*^, *myc*^*P0*^, and *bam*>*myc*^*OE*^ ovaries. Germaria are outlined with dotted lines. Reduction in either IIS or Myc depletes *InR* mRNA in the germarium, while *myc* over-expression in region 2A ectopically induces *InR* mRNA. (B) Quantification of *InR* mRNA intensity from cysts in germarium region 2A and 2B of ovaries with indicated genotypes. Intensities are normalized to the value of *ctrl* at region 2B. (C) Germaria from *nos*>*ctrl, nos*>*chico*^*RNAi*^, *myc*^*P0*^, and *bam*>*myc*^*OE*^ ovaries stained with anti-p-AKT and anti-1B1. Decrease in either IIS or Myc reduces IIS activity in the germarium, while *myc* over-expression in region 2A ectopically induces IIS activity. (D) Quantification of p-AKT intensity in region 2A and 2B cysts of ovaries with indicated genotypes. Intensities are normalized to the value of *ctrl* at region 2B. Error bars represent SEM. *p < 0.005.

### The JNK-IIS-Myc relay is essential for female fertility and mtDNA selective inheritance

So far we have established that the JNK-IIS-Myc relay is critical for ETC activity, mtDNA expression, replication, and inheritance in the ovary. Next, we explored its physiological impact on reproduction. While both *chico* mutant females and those with germline clones of *bsk*^*1*^ produced similar amount of eggs as control, they failed to generate adequate amount of mtDNA to deposit in eggs (Fig 7A). Thus, their eggs had significantly reduced mtDNA level and hatching rates (Fig 7B).

**Figure 7.**
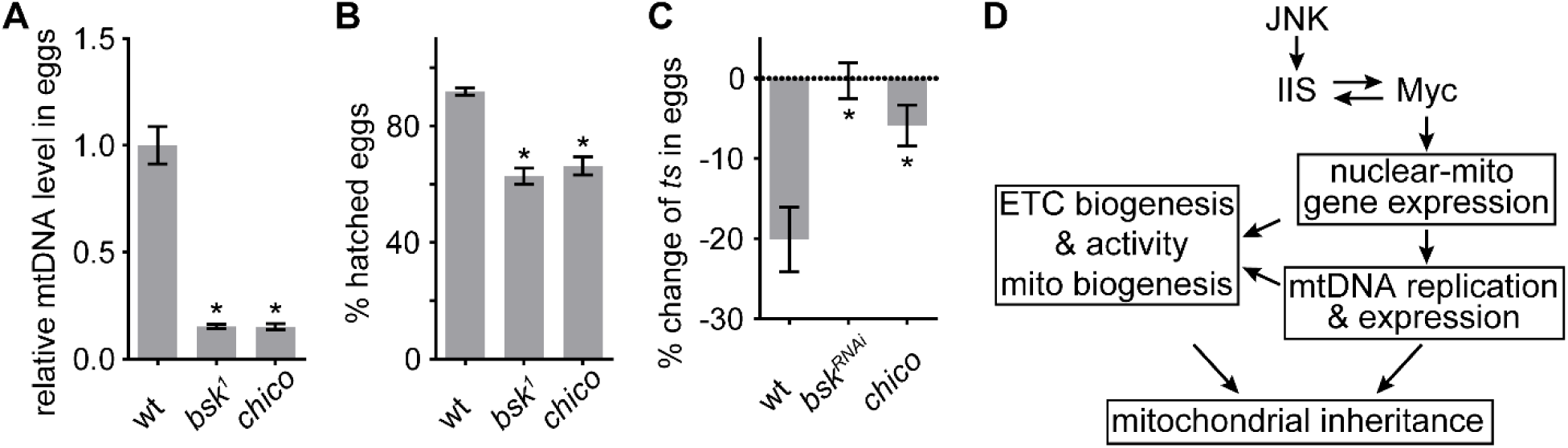
JNK-IIS relay is essential for mtDNA selective inheritance and fertility. (A) Quantification of relative mtDNA content in eggs produced by mothers of either GLC for *bsk*^*1*^ or *chico*^*1*/*KG*^ mutant. Relative mtDNA levels are determined by qPCR for *mt:CoI* and *his4* copies, and normalized to the level in wt. (B) Hatching rates of eggs produced by female flies of either GLC for *bsk*^*1*^ or *chico*^*1*/*KG*^mutant are significantly reduced compared to the hatching rates of wt eggs. (C) Quantification of *mt:Col*^*T300I*^ transmission from females with indicated nuclear genotypes. In wt females, the mtDNA mutation is counter-selected, resulting in ∼20% fewer mutant mtDNAs in the progeny than in the mothers. This counter-selection is diminished in *chico*^*1IKG*^ mutant or *nos*>*bsk*^*RNAi*^ mothers. Mother flies heteroplasmic for *mt:Col*^*T300I*^ were cultured at 29°C. (D) Schematics of the developmental signaling relay initiated from late germarium that primes mitochondrial respiration, and mtDNA replication and inheritance. Error bars represent SEM. *p < 0.005.

During oogenesis, the wild-type mitochondrial genomes enriched in healthy mitochondria out-compete lethally mutated mtDNA afflicting defective mitochondria (Hill et al., 2014). Thus, we asked whether abolishing IIS and JNK signaling would impair mtDNA selective inheritance in heteroplasmic females harboring both wt and a temperature-sensitive lethal mutation, *mt:CoI*^*T300I*^ (Hill et al., 2014). Consistent with previous studies, eggs contain ∼20% less *mt:CoI*^*T300I*^ mtDNA on average than their mothers at a restrictive temperature (Fig 7C). However, down-regulation of either IIS or JNK signaling abolished the counter-selection of *mt:CoI*^*T300I*^ genome (Fig 7C). Together, these results stress that although JNK is transiently activated in the late germarium, it triggers a developmental signaling relay that has profound impacts on mitochondrial inheritance through activation of ETC biogenesis and mtDNA replication.

## DISCUSSION

mtDNA replication in the *Drosophila* ovary relies on active respiration (Hill et al., 2014), suggesting that ETC activity might be subject to the same spatial-temporal regulation as mtDNA replication. In this study, we address this question and further elucidate the underlying developmental regulations of ETC activity and mtDNA biogenesis in ovary. We reveal that ETC complexes, exemplified by COX/SDH are inactive in the GSCs and dividing cysts from germarium region 1 to 2A, but sharply activated in region 2B, and remain active until stage 10 follicles, in which a metabolic reprograming takes place (Sieber et al., 2016). This spatial pattern mirrors that of mtDNA replication in the *Drosophila* ovary, further substantiating an essential role of mitochondrial respiration in mtDNA inheritance, both quantitively and qualitatively. We also demonstrate that the ETC activation is accompanied with an upregulation of the expression of ETC genes on both nuclear and mitochondrial genomes. Interestingly, MDI that drives the local translation of nuclear encoded mitochondrial proteins on the mitochondrial outer membrane and TFAM that governs mtDNA replication and transcription (Falkenberg et al., 2007; Zhang et al., 2016), exhibit the same developmental pattern as mitochondrial respiration in the germarium. Collectively, these proteins would boost the biogenesis of ETC in region 2B germarium and growing egg chambers. In an ovariole, different stages of developing germ cells reside in the same microenvironment and experience same oxygen tension. Thus, the mitochondrial respiratory activity is likely to be determined by the abundance of ETC, which is controlled by transcriptional activation.

To understand how ETC biogenesis is regulated, we conducted a candidate RNAi screening for genes that are required for boosting COX/SDH activities in ovary. Myc emerged as one of the strongest hits, and a hypermorphic allele of *myc*^*P0*^ largely abolished ETC activities and mtDNA replication in the germarium. Moreover, the spatial pattern of Myc protein mirrors mtDNA replication and ETC activity, further supporting its essential role in transcriptional activation of ETC biogenesis. RNA sequencing data demonstrate that Myc broadly stimulates the gene expression in the *Drosophila* ovary, including many nuclear encoded ETC genes and factors required for mtDNA replication and expression. Our results are consistent with studies on mammal cultured cells showing that Myc can stimulate mitochondrial genes expression and enhance mitochondrial respiration (Li et al., 2005; Stine et al., 2015). Myc overexpression in different cell types or different conditions, sometimes give rise to different transcriptional output (de la Cova et al., 2014). These observations underscore that Myc family proteins often associate with other cofactors and exert a broad and complex transcriptional activation or repression in a cell/tissue specific manner (Cowling and Cole, 2006; Hann, 2014). We also found that 130 transcription regulators, including *Srl* (fly homolog of human PGC-1) and CG32343 (fly homolog of GABPB2), were affected in the *myc*^*P0*^ ovary. PGC-1 proteins belong to an evolutionarily conserved family that integrates mitochondrial biogenesis and energy metabolism with a variety of cellular processes (Lin et al., 2005). In *Drosophila, Srl* regulates the expression of a subset of nuclear encoded mitochondrial genes (Tiefenbock et al., 2010). Mammalian GABPB2 is a regulatory subunit of the Nuclear Respiratory Factor 2 complex that regulates the expression of a small set of nuclear encoded mitochondrial proteins (Kelly and Scarpulla, 2004). Therefore, additional tiers of transcriptional regulations downstream of Myc are likely involved to boost ETC biogenesis.

While *myc* transcription is uniform in the germarium, Myc protein is elevated at region 2B and remained at high level until the stage 10 egg chamber, highlighting that Myc protein level is mainly regulated *via* post transcriptional mechanisms,. IIS and JNK also emerged from our candidate screening, and both were further confirmed to be required for triggering ETC biogenesis and mtDNA replication. We found that IIS activity, marked by both p-AKT and p-GSK3 staining, displayed a similar pattern to that of Myc. Additionally, elevated IIS activity was required to establish a high level of Myc and to activate ETC in late germarium stage. Our result is line with a previous study showing that decreased IIS activity relieves the inhibition on GSK3, which leads to mitochondrial quiescence at later stages of oogenesis (Sieber et al., 2016). Importantly, our work uncovers Myc as the downstream effector of IIS on the regulation of respiration and mtDNA biogenesis in the ovary.

We noticed that *InR* transcription was down-regulated in the *myc* mutant ovary, suggesting a positive feedback regulation between IIS and Myc. This regulatory loop maintains high levels of both Myc protein and IIS in the mid-stage follicles, where prodigious mitochondrial biogenesis and massive cell growth take place. However, it does not explain how this loop is activated in the first place at the late germarium stages. We found that JNK was transiently activated in germ cells at the germarium 2B, but decreased in budding egg chambers and sharply diminished thereafter. High level and sustained JNK activity often leads to apoptosis. However, cell death is rarely observed in germaria of flies cultured under the normal condition. Thus, temporal JNK activation at late germarium must be associated with other cellular processes. We reveal that transiently-elevated JNK activity is sufficient to increase *InR* mRNA level, which in-turn boosts IIS and stabilizes Myc protein. Currently, the potential link between JNK and IIS is well-understood. The metastatic *Drosophila* epithelium cells require JNK to up-regulate *InR* expression through wingless signaling to promote cell survival and proliferation (Hirabayashi et al., 2013). However, no genes in wingless signaling emerged from our RNAi screening in germ cells. Molecular mechanisms that links JNK activation to *InR* expression in ovary remains to be explored.

JNK-dependent transcriptional program can be activated by various cellular stresses and cell-cell signaling (Rios-Barrera and Riesgo-Escovar, 2013). In region 2B germarium, the follicle cells extend and migrate laterally across the germarium to wrap around the 16-cells cyst (Nystul and Spradling, 2007). Thus, follicle cells may active JNK in germ cells through paracrine signals, such as TNF-α. Alternatively, the process of follicle cells enveloping and compressing the 16-cell cyst may generate mechanical stresses that subsequently activates JNK. Regardless, our work uncovers a novel function of JNK in energy metabolism and mitochondrial biogenesis besides its well established roles in controlling cell apoptosis, growth, and proliferation.

Studies in a variety of animal models have showed that reproductive aging of females is tightly associated with decreased IIS (Templeman and Murphy, 2018). Interestingly, oocytes of aged females often have higher incidence of mtDNA lesions and lower mtDNA copy number (Chan et al., 2005). Thus, our findings on developmental control of mitochondrial biogenesis and mtDNA replication via IIS may be a conserved mechanism in metazoans. Our studies could be a molecular framework to further understand the control of mitochondrial biogenesis and mtDNA inheritance in animals.

## STAR METHODS

### Detailed methods are provided in the online version of this paper and include the following

- **KEY RESOURCES TABLE**
- **CONTACT FOR REAGENT AND RESOURCE SHARING**
- **EXPERIMENTAL MODEL AND SUBJECT DETAILS**
- **METHODS DETAILS**
- **QUANTIFICATION AND STATISTICAL ANALYSIS**
- **DATA AVAILABILITY**

## SUPPLEMENTAL INFORMATION

Supplemental Information includes seven figures and three tables.

## ACKNOWLEDGEMENTS

We thank F Chanut for comments and editing on the manuscript; D. Drummond-Barbosa, E. Geisbrecht, L. Johnston, E. Moreno, O. Schuldiner, Bloomington *Drosophila* Stock Center, and Vienna Drosophila Resource Center for various fly lines; NHLBI DNA Sequencing and Genomics Core, Bioinformatics and Computational Biology Core and, Bestgene Inc. for technical assistance. Funding: This work is supported by NHLBI Intramural Research Program.

## AUTHOR CONTRIBUTIONS

Z-H.W. and H.X. conceived the project and design the experiments. Z-H.W. and Y.L. performed the experiments. Z-H.W., V.C., M.P. and H. X. analyzed data. Z-H.W. and H.X. wrote the paper.

## SUPPLEMENTAL INFORMATION

## SUPPLEMENTAL TABLES

Table S1 Candidate RNAi screen.

Table S2 Differential expression of mRNAs in *myc*^*P0*^ mutant ovaries compared with wt ovaries.

Table S3 Sequences of FISH probes.

## STAR METHODS

## KEY RESOURCES TABLE

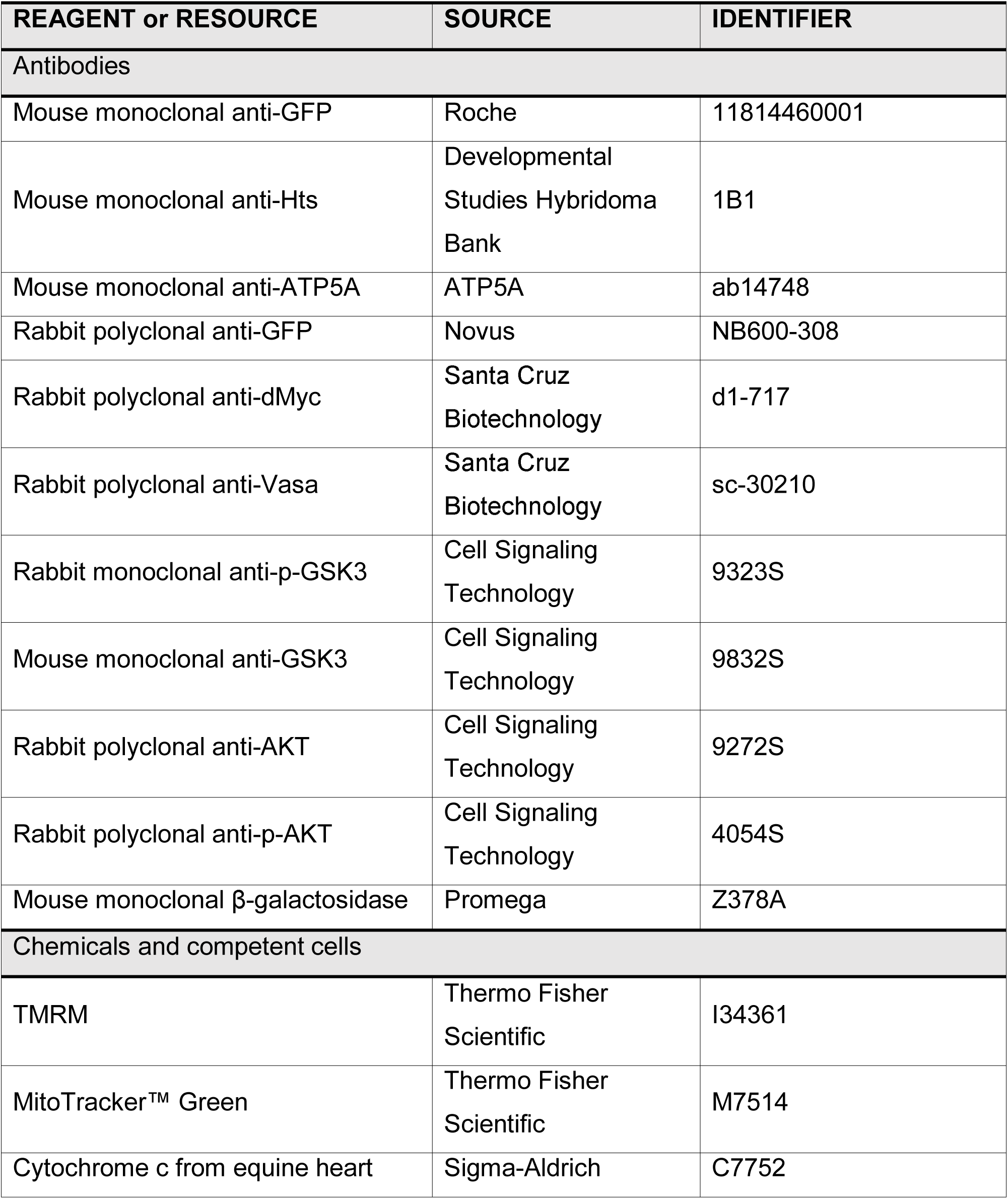
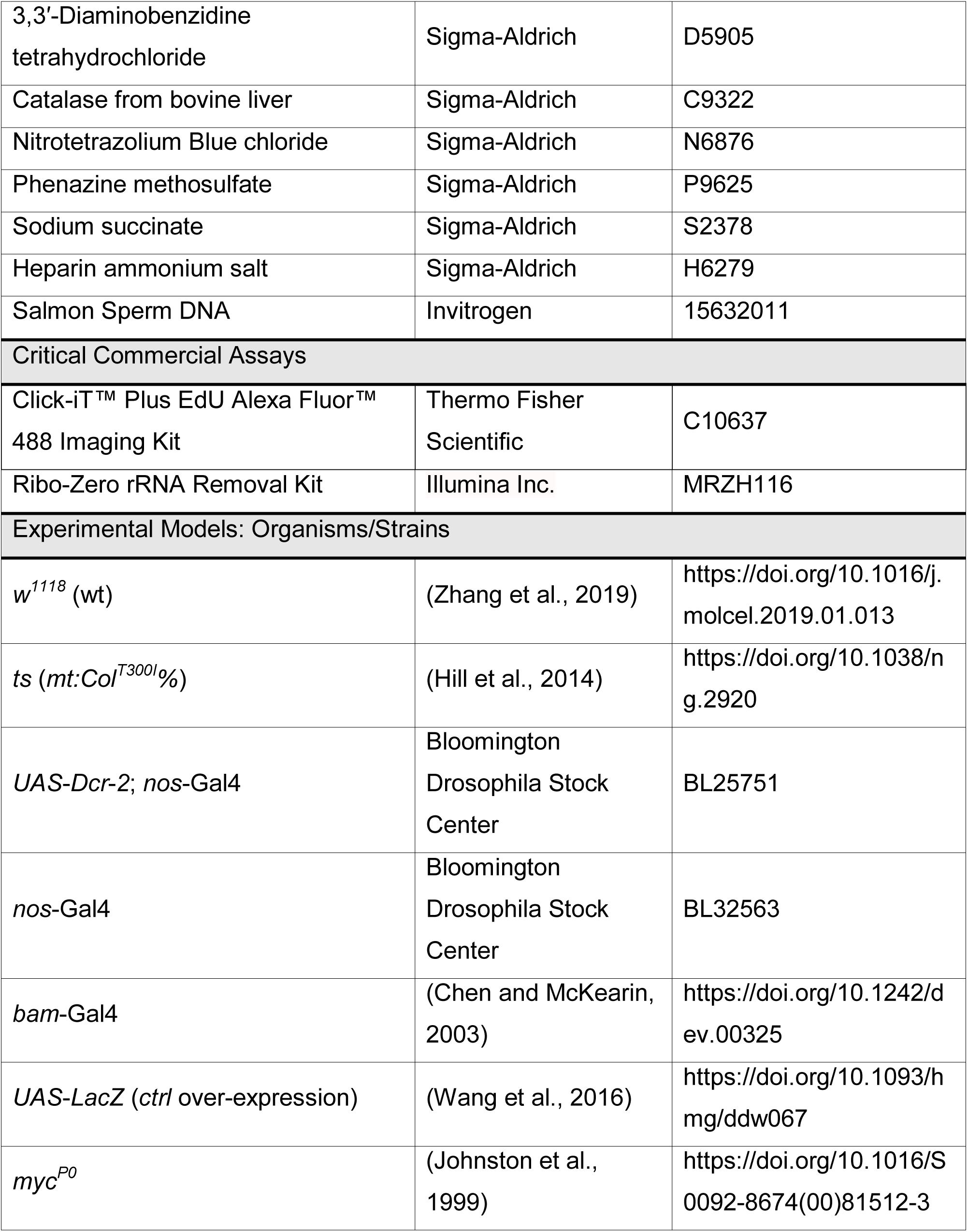
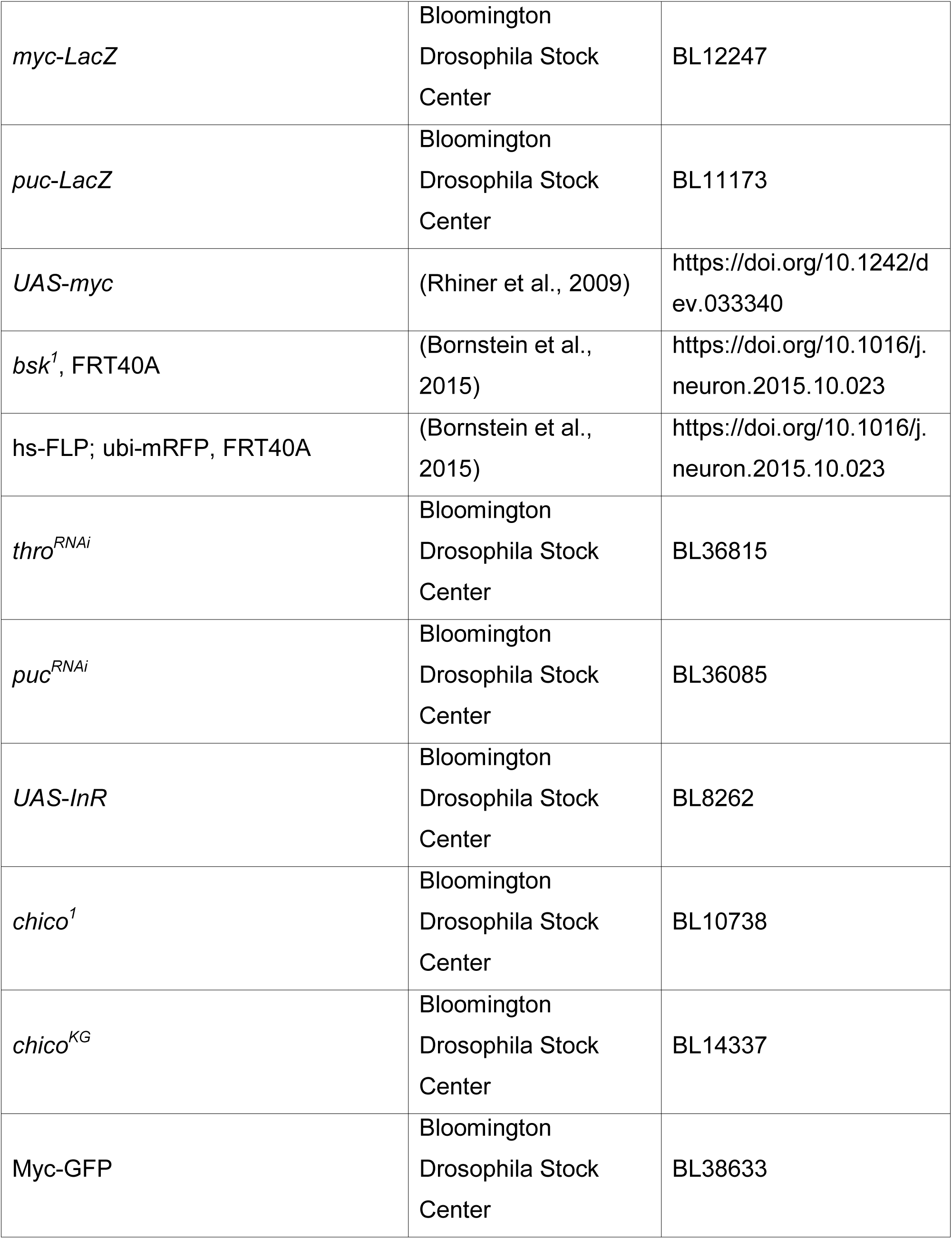
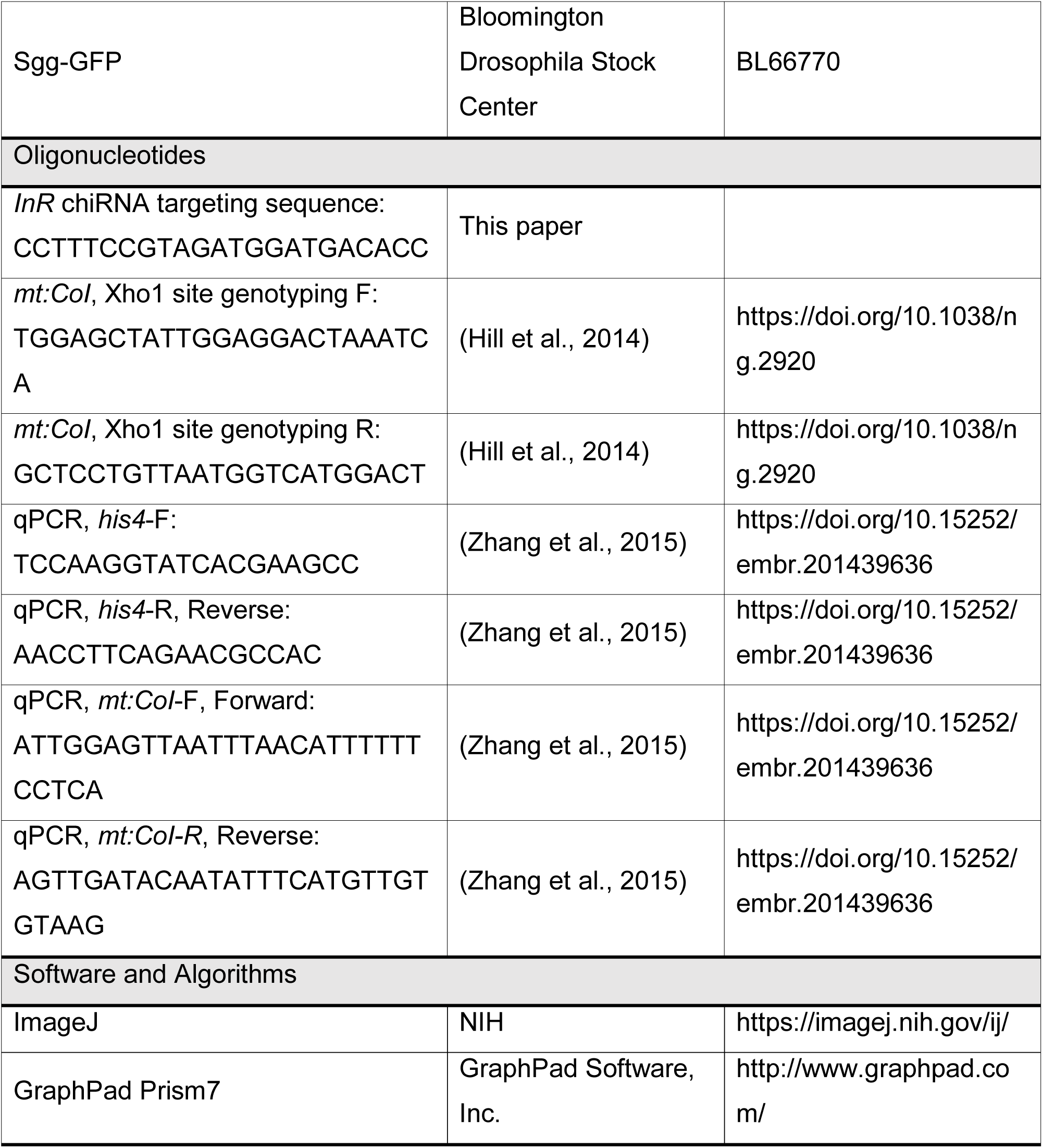

## CONTACT FOR REAGENT AND RESOURCE SHARING

Further information and requests for resources and reagents should be directed to and will be fulfilled by the Lead Contact, Hong Xu.

## EXPERIMENTAL MODEL AND SUBJECT DETAILS

### Drosophila Genetics

Flies were maintained on standard BDSC cornmeal medium at 25°C. RNAi lines obtained from either the Bloomington Stock Center or the Vienna Drosophila RNAi Center are listed in KEY RESOURCES TABLE. Embryo hatch assay was performed as previously described (Zhang et al., 2019).

## METHOD DETAILS

### Measurement of mtDNA copy number and quantification of heteroplasmy

Total DNA was isolated from eggs with QIAamp DNA Micro Kit (Qiagen). The mtDNA copy number was measured by quantitative real-time PCR with primers targeting to cytochrome c oxidase subunit I (mt:CoI) and His4 genes. Quantification of heteroplasmy was performed as described previously (Zhang et al., 2019). Heteroplasmic female flies were transferred from 18 °C to 29 °C after eclosion. Each female was mated with 5 wt males. Ten eggs produced from the day 7 at 29 °C were collected. The genomic DNA from female flies and eggs was extracted and their heteroplasmy levels were determined as shown before (Hill et al., 2014).

### CRISPR/Cas9 in flies

To tag InR with EGFP into its endogenous locus, a DNA fragment comprising 1kb upstream of *InR* stop codon, EGFP, a fragment containing GMR-Hid flanked by FRT sequences, and 1kb downstream of *InR* stop codon was inserted into a pOT2 vector. This donor construct and a chiRNA construct inserted with a targeting sequence CCTTTCCGTAGATGGATGACACC were injected into the embryos of M{vas-Cas9}ZH-2A (BL51323) by Bestgene Inc.. G1 adults were screened for loss of eye phenotype. Insertion events were further confirmed by PCR using two pairs of primers: 1, InR-GenomeF1: ATGATGTCATCGGTGGGTCCTCAC, EGFP-seqR: CTTGTAGTTGCCGTCGTCCTTGAA; 2, InR-GenomeF2: AGCACATTGTGTCAGTCTTCG, InR-GenomeR: CTCATTTTCCGAAGCTTGGCTTCC.

### RNA sequencing and RNA-seq analysis

Total RNA was extracted by Trizol (Life Technologies) following its standard protocol. Poly (A) capture libraries were generated at the DNA Sequencing and Genomics Core, NHLBI, NIH. RNA sequencing was performed with using an Hiseq3000 (Illumina) and 75-bp pair-end reads were generated at the DNA Sequencing and Genomics Core, NHLBI, NIH. Raw sequence reads were quality-trimmed using Trim Galore! (v0.3.7) and aligned using Tophat2 (v2.0.14) against the Dm6 reference genome. Uniquely mapped paired-end reads were then used for subsequent analyses. FeatureCounts was used for gene level abundance estimation. Principal component analysis (PCA) was used to assess outlier samples. Genes were kept in the analysis if they had Counts Per Million (CPM>1) in at least half the samples. We adjusted for multiple testing by reporting the FDR q-values for each feature. Features with q < 5% were declared as genome-wide significant. Genes with 3 or more fold changes on mRNA level in *myc*^*P0*^ mutant compared with wt were considered differentially expressed. Gene Ontology (GO) was used to analyze gene set enrichment. FDR q-values were estimated to correct the p-values for the multiple testing issue.

### Live image

Live image of fly ovaries was performed as previously reported (Zhang et al., 2019). Ovaries from wt flies were stained with TMRM and MTgreen for 20 min followed by PBS washes for 3 times. Ovaries from Sgg-GFP flies were dissected. Ovarioles were isolated and immerged in halocarbon oil on coverslips, then live imaged with a Perkin Elmer Ultraview system. The ratiometric image was generated with ImageJ, which the intensity of red channel is divided by that of green channel.

### Immunohistochemistry and fluorescence RNA in situ hybridization (FISH)

Ovary dissection, immunostaining and EdU incorporation were performed as described before (Hill et al., 2014). Stellaris FISH probes against, *Cyt*-*C1, Cyt*-*B, CoxIV, CoxIII*, or *InR* mRNA were synthesized from Bioserch Technologies. Sequences of the probes are shown in table S3. FISH of *Drosophila* ovaries was conducted as previously described (Trcek et al., 2017). Confocal images were collected by a Perkin Elmer Ultraview system or Instant Sim (iSIM) Super-Resolution Microscope. All images were processed with Image J.

## QUANTIFICATION AND STATISTICAL ANALYSIS

All statistical analyses were conducted with Prism 7 (GraphPad Software). Error bars in all charts represent standard errors. P values were performed with Two-tailed Student’s t test. Statistical significance of difference was considered when P < 0.05.

## DATA AVAILABILITY

The data were deposited in Gene Expression Omnibus of NCBI (Edgar et al., 2002) and will be available with accession number (GEO: GSE126997).

## SUPPLEMENTAL FIGURES AND FIGURE LEGENDS

**Figure S1.**
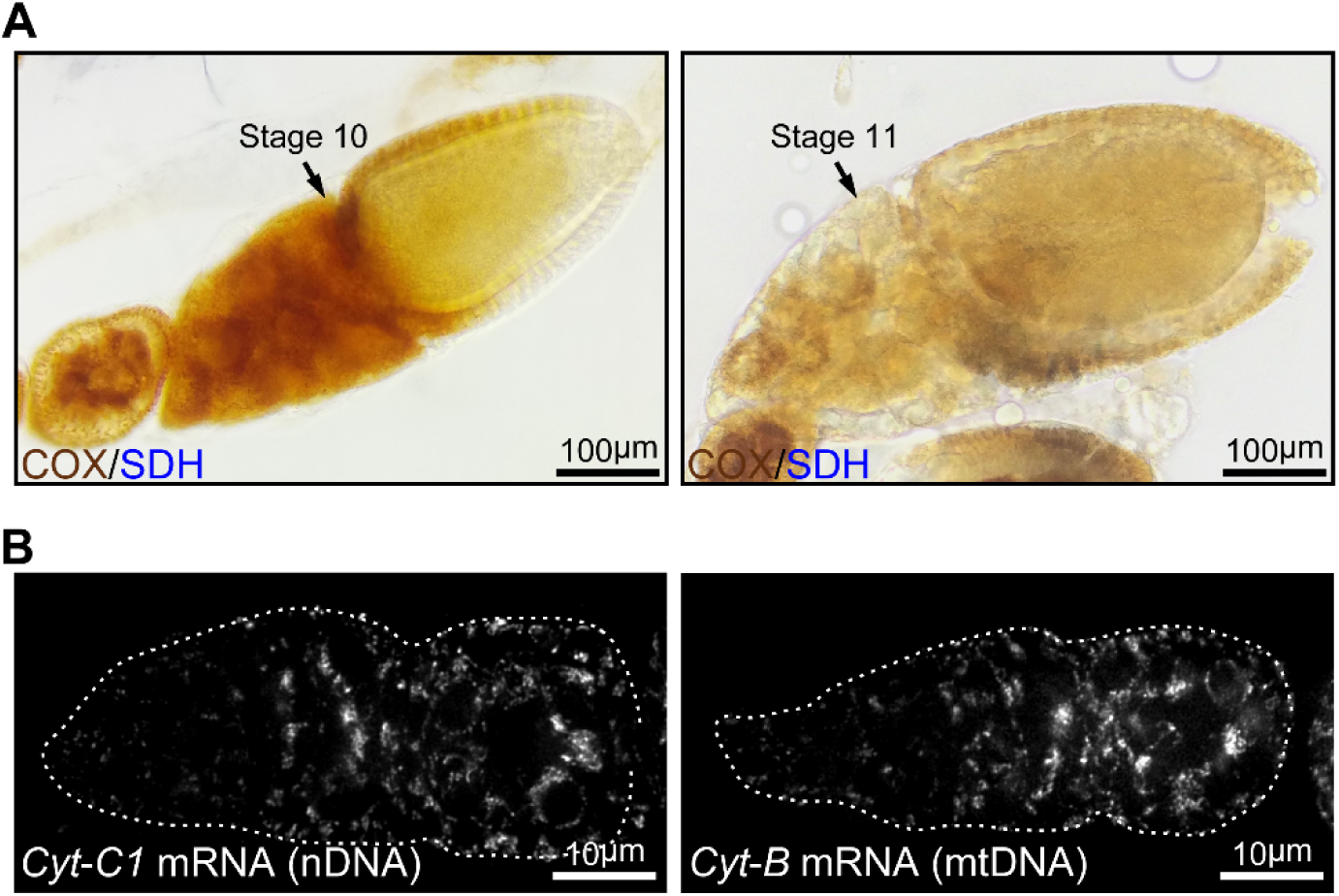
ETC activity at middle stages of oogenesis and expression of complex III genes in the germarium. (Related to Figure 1) (A) Egg chambers from wt flies stained with COX/SDH. Note that ETC activity is high in a stage 10 egg chamber, while it is dramatically reduced in a stage 11 egg chamber. (B) Visualization of the *Cyt*-*C1* and *Cyt*-*B* mRNA in germaria from wt flies with fluorescently labelled DNA probes by FISH. Germaria are outlined with dotted lines.

**Figure S2.**
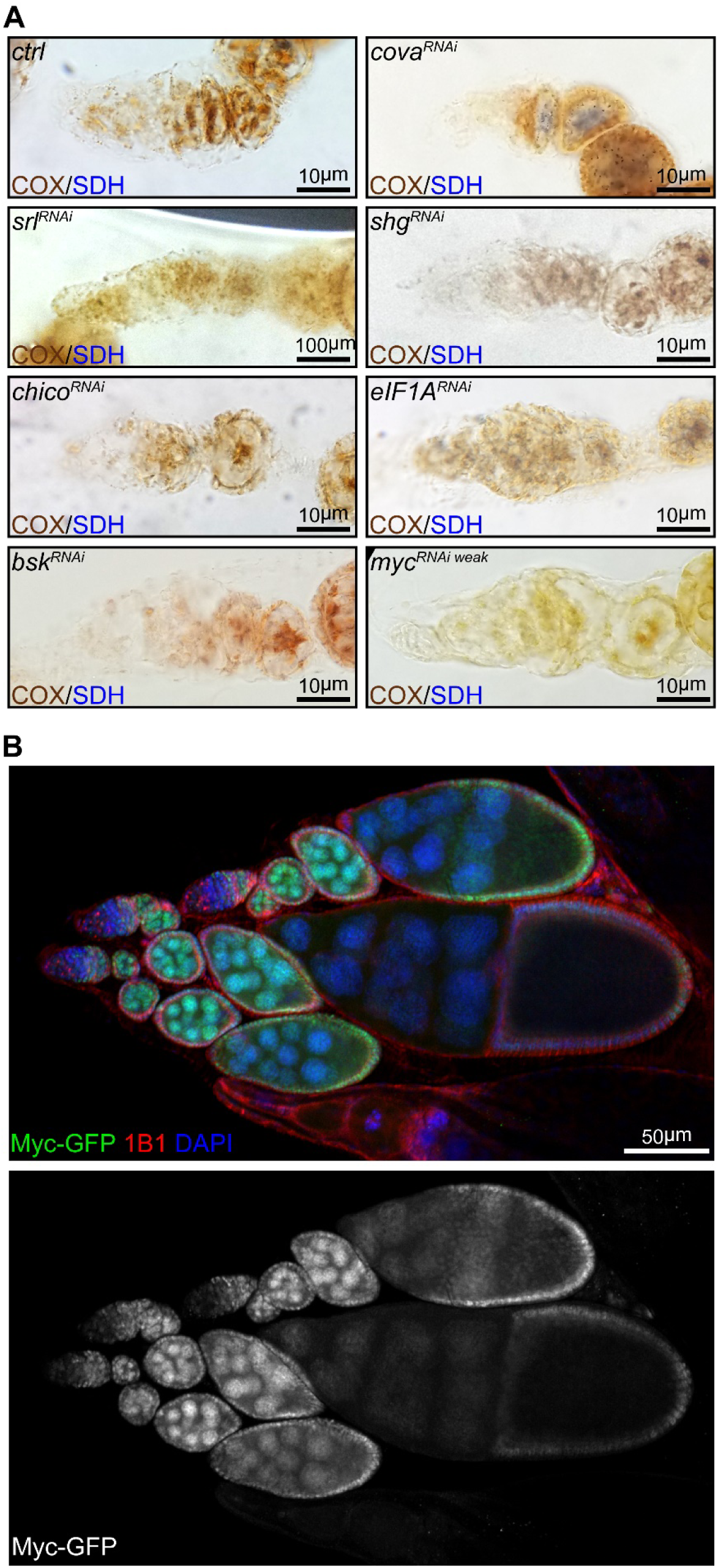
ETC activity in representative candidate RNAi screen and Myc protein pattern in the ovary. (Related to Figure 2) (A) COX/SDH staining for germaria with reduced ETC activity (positive hits) from the candidate RNAi screen. (B) A representative low magnification image of ovarioles from flies endogenously expressing Myc-GFP stained with anti-GFP, anti-1B1, and DAPI. Note that Myc level is low in early germarium stages, becomes high from germarium region 2B, and reduces from the stage 10 egg chamber.

**Figure S3.**
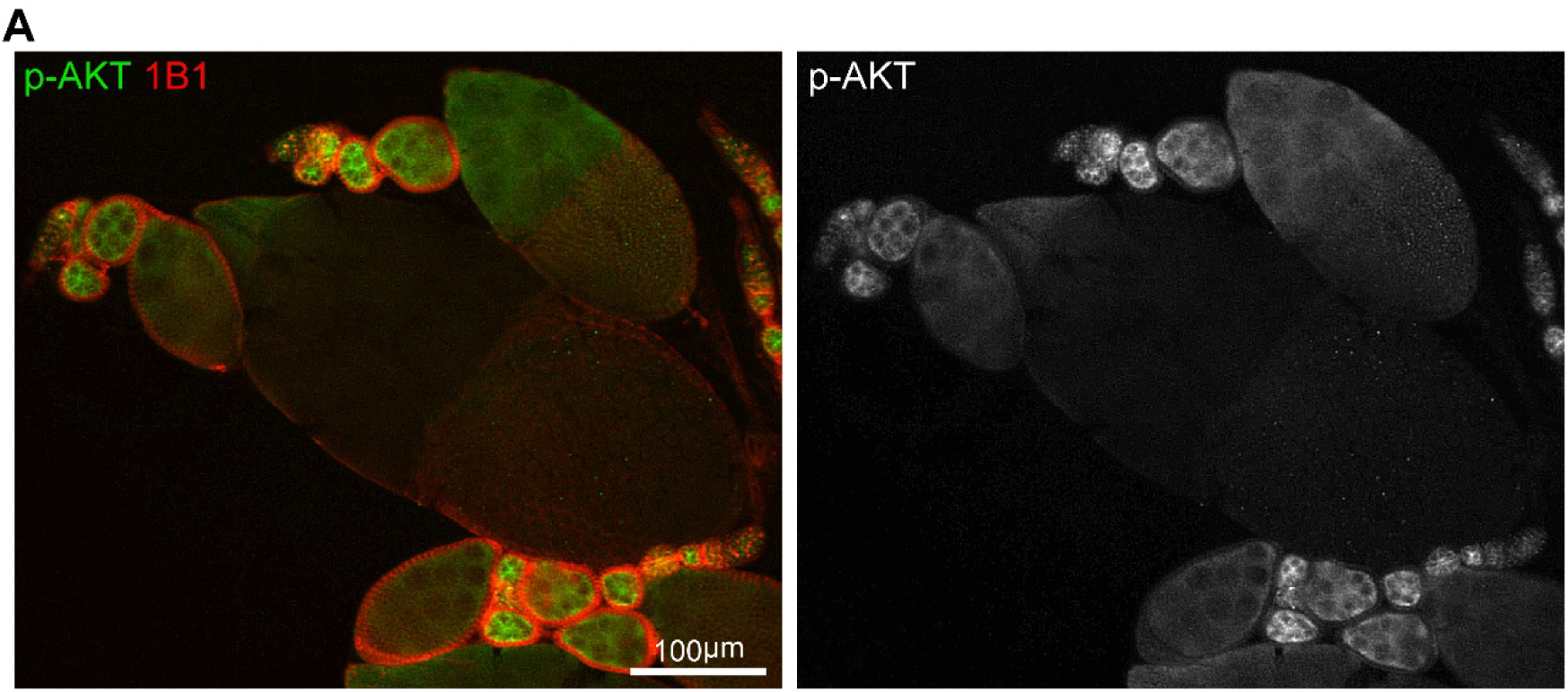
IIS activity in the ovary. (Related to Figure 3) (A) A low magnification image of ovarioles stained with anti-p-AKT and anti-1B1. Note that IIS activity is low in early germarium stages, becomes high from late germarium, and decreases from the stage 10 egg chamber, the same pattern as Myc protein.

**Figure S4.**
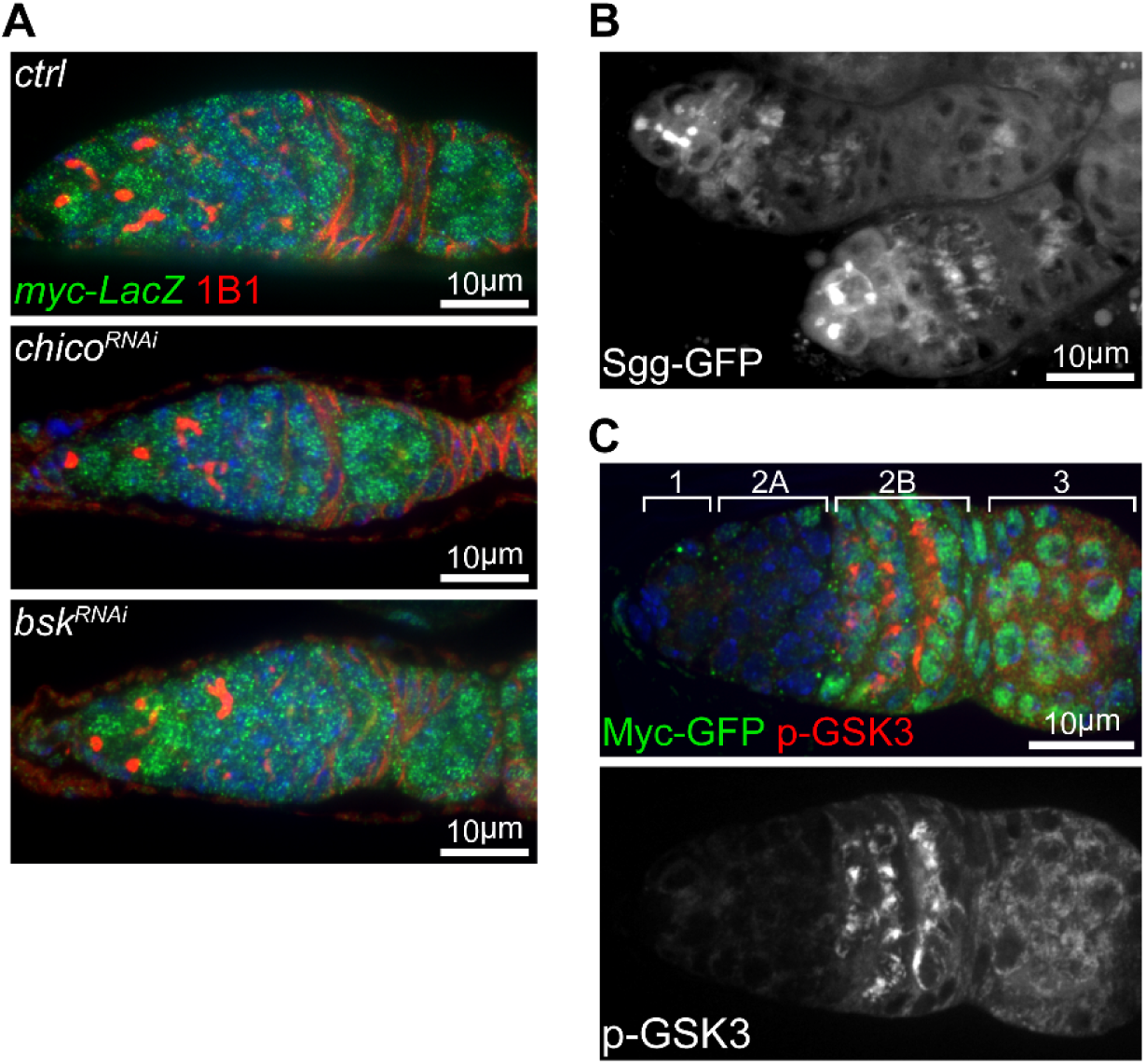
Myc is post-transcriptionally regulated by the IIS-GSK3 cascade. (Related to Figure 4) (A) Germaria from *nos*>*ctrl, nos*>*chico*^*RNAi*^, and *nos*>*bsk*^*RNAi*^ ovaries expressing LacZ under the control of *myc* endogenous promoter stained with anti-β-galactosidase and anti-1B1. (B) A representative live imaging of germaria from flies expressing GFP-tagged Sgg under its endogenous promoter. (C) A germarium from wt flies endogenously expressing Myc-GFP stained with anti-GFP (green) and anti-p-GSK3 (red). Germarium regions are indicated. Myc protein highly corresponds to inhibited form of GSK3.

**Fig S5.**
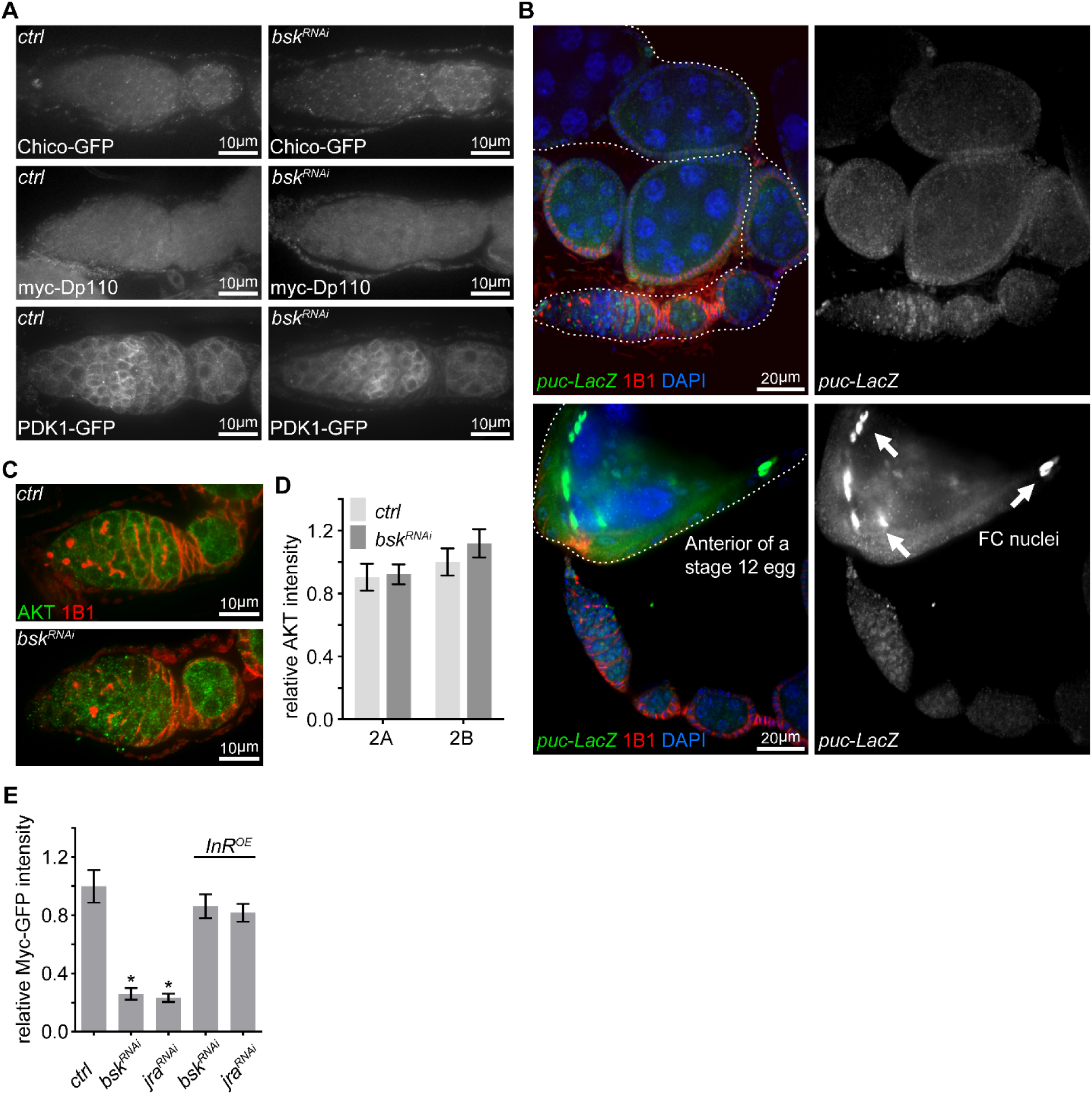
Other IIS components are not altered by *bsk* RNAi. (Related to Figure 5) (A) Confocal images for germaria expressing Chico-GFP, myc-Dp110 or PDK1-GFP in the background of *ctrl* or *bsk*^*RNAi*^ from ovaries stained with anti-GFP or anti-myc. (B) Ovaries expressing LacZ driven by the *puc* promoter stained with anti-β-galactosidase, anti-1B1, and DAPI. Note that JNK activity is moderately induced in late germarium and decreased in the following stages, while JNK activity is much stronger in follicle cells of a maturing egg. (C) Germaria from *nos*>*ctrl* and *nos*>*bsk*^*RNAi*^ flies stained with anti-AKT and anti-1B1. (D) Quantification of AKT intensity from cysts in germarium region 2A and 2B of ovaries with indicated genotypes. Intensities are normalized to the value of *ctrl* at region 2B. (E) Quantification of relative Myc-GFP intensity in region 2B cysts of ovaries illustrated in Fig 5H. Error bars represent SEM. *p < 0.005.

**Figure S6.**
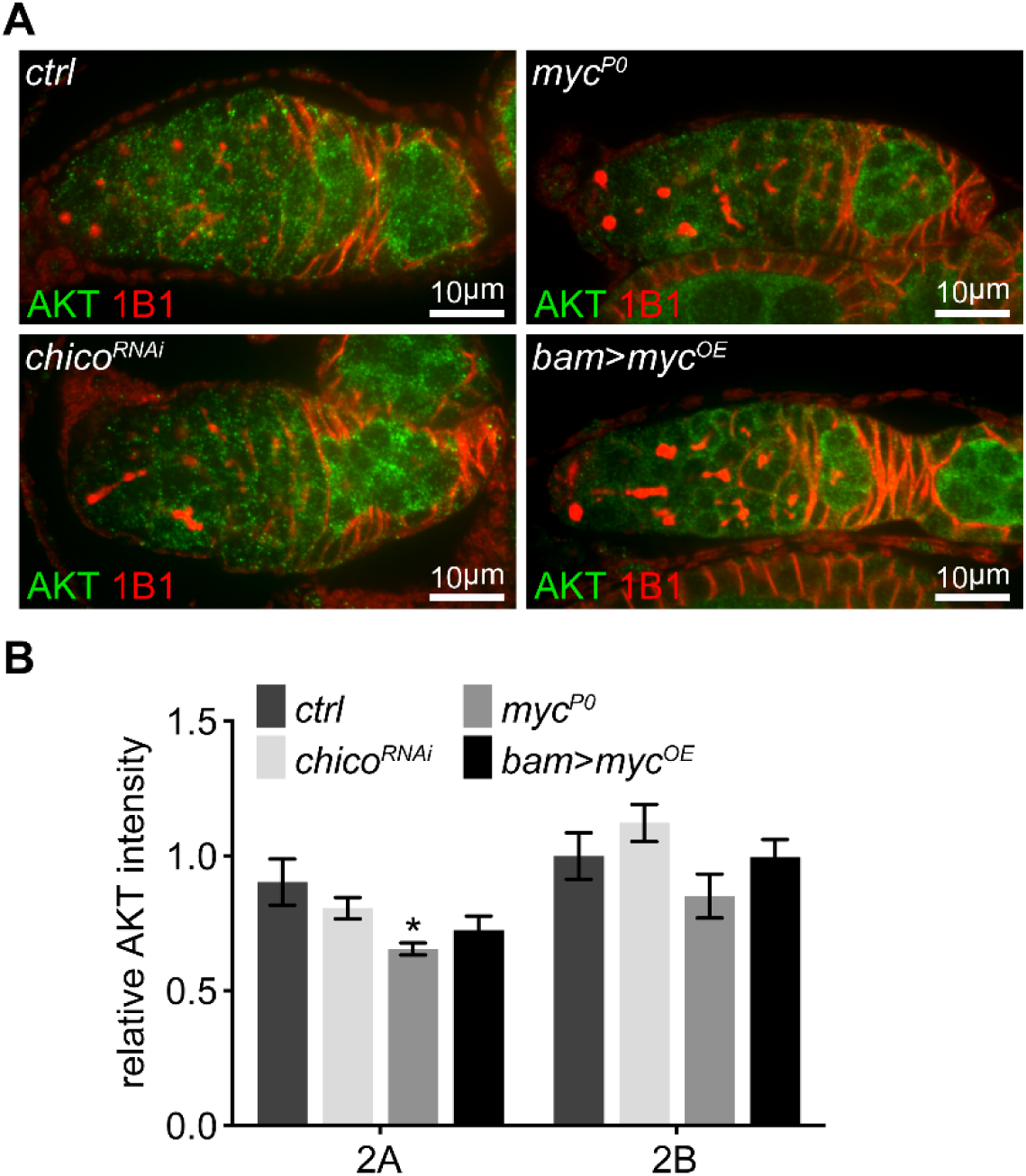
AKT level is normal when IIS is decreased, but slightly reduced in the *myc*^*P0*^ mutant. (Related to Figure 6) (A) Germaria from *nos*>*ctrl, nos*>*chico*^*RNAi*^, *myc*^*P0*^, and *bam*>*myc*^*OE*^ ovaries stained with anti-AKT and anti-1B1. (B) Quantification of AKT intensity in region 2A and 2B from ovaries with indicated genotypes. Error bars represent SEM. *p < 0.005.

## REFERENCES

Basson, M.A. (2012). Signaling in cell differentiation and morphogenesis. Cold Spring Harb Perspect Biol 4.

Bohni, R., Riesgo-Escovar, J., Oldham, S., Brogiolo, W., Stocker, H., Andruss, B.F., Beckingham, K., and Hafen, E. (1999). Autonomous control of cell and organ size by CHICO, a Drosophila homolog of vertebrate IRS1-4. Cell 97, 865–875.

Chan, C.C., Liu, V.W., Lau, E.Y., Yeung, W.S., Ng, E.H., and Ho, P.C. (2005). Mitochondrial DNA content and 4977 bp deletion in unfertilized oocytes. Mol Hum Reprod 11, 843–846.

Claveria, C., and Torres, M. (2016). Cell Competition: Mechanisms and Physiological Roles. Annu Rev Cell Dev Biol 32, 411–439.

Cowling, V.H., and Cole, M.D. (2006). Mechanism of transcriptional activation by the Myc oncoproteins. Semin Cancer Biol 16, 242–252.

de la Cova, C., Senoo-Matsuda, N., Ziosi, M., Wu, D.C., Bellosta, P., Quinzii, C.M., and Johnston, L.A. (2014). Supercompetitor status of Drosophila Myc cells requires p53 as a fitness sensor to reprogram metabolism and promote viability. Cell Metab 19, 470-483.

Desvergne, B., Michalik, L., and Wahli, W. (2006). Transcriptional regulation of metabolism. Physiol Rev 86, 465–514.

Falkenberg, M., Larsson, N.G., and Gustafsson, C.M. (2007). DNA replication and transcription in mammalian mitochondria. Annu Rev Biochem 76, 679–699.

Gallant, P. (2013). Myc function in Drosophila. Cold Spring Harb Perspect Med 3, a014324.

Garofalo, R.S. (2002). Genetic analysis of insulin signaling in Drosophila. Trends Endocrinol Metab 13, 156–162.

Greer, C., Lee, M., Westerhof, M., Milholland, B., Spokony, R., Vijg, J., and Secombe, J. (2013). Myc-dependent genome instability and lifespan in Drosophila. PLoS One 8, e74641.

Hann, S.R. (2014). MYC cofactors: molecular switches controlling diverse biological outcomes. Cold Spring Harb Perspect Med 4, a014399.

Hill, J.H., Chen, Z., and Xu, H. (2014). Selective propagation of functional mitochondrial DNA during oogenesis restricts the transmission of a deleterious mitochondrial variant. Nat Genet 46, 389–392.

Hirabayashi, S., Baranski, T.J., and Cagan, R.L. (2013). Transformed Drosophila cells evade diet-mediated insulin resistance through wingless signaling. Cell 154, 664–675.

Johnston, L.A., Prober, D.A., Edgar, B.A., Eisenman, R.N., and Gallant, P. (1999). Drosophila myc regulates cellular growth during development. Cell 98, 779–790.

Kelly, D.P., and Scarpulla, R.C. (2004). Transcriptional regulatory circuits controlling mitochondrial biogenesis and function. Genes Dev 18, 357–368.

LaFever, L., and Drummond-Barbosa, D. (2005). Direct control of germline stem cell division and cyst growth by neural insulin in Drosophila. Science 309, 1071–1073.

Li, F., Wang, Y., Zeller, K.I., Potter, J.J., Wonsey, D.R., O’Donnell, K.A., Kim, J.W., Yustein, J.T., Lee, L.A., and Dang, C.V. (2005). Myc stimulates nuclearly encoded mitochondrial genes and mitochondrial biogenesis. Mol Cell Biol 25, 6225–6234.

Lin, J., Handschin, C., and Spiegelman, B.M. (2005). Metabolic control through the PGC-1 family of transcription coactivators. Cell Metab 1, 361–370.

Martin-Blanco, E., Gampel, A., Ring, J., Virdee, K., Kirov, N., Tolkovsky, A.M., and Martinez-Arias, A. (1998). puckered encodes a phosphatase that mediates a feedback loop regulating JNK activity during dorsal closure in Drosophila. Genes Dev 12, 557–570.

Maurer, U., Preiss, F., Brauns-Schubert, P., Schlicher, L., and Charvet, C. (2014). GSK-3 - at the crossroads of cell death and survival. J Cell Sci 127, 1369–1378.

May-Panloup, P., Chretien, M.F., Malthiery, Y., and Reynier, P. (2007). Mitochondrial DNA in the oocyte and the developing embryo. Curr Top Dev Biol 77, 51–83.

Nagarkar-Jaiswal, S., Lee, P.T., Campbell, M.E., Chen, K., Anguiano-Zarate, S., Gutierrez, M.C., Busby, T., Lin, W.W., He, Y., Schulze, K.L., et al. (2015). A library of MiMICs allows tagging of genes and reversible, spatial and temporal knockdown of proteins in Drosophila. Elife 4.

Neto-Silva, R.M., de Beco, S., and Johnston, L.A. (2010). Evidence for a Growth-Stabilizing Regulatory Feedback Mechanism between Myc and Yorkie, the Drosophila Homolog of Yap. Dev Cell 19, 507–520.

Nystul, T., and Spradling, A. (2007). An epithelial niche in the Drosophila ovary undergoes long-range stem cell replacement. Cell Stem Cell 1, 277–285.

Oldham, S., and Hafen, E. (2003). Insulin/IGF and target of rapamycin signaling: a TOR de force in growth control. Trends Cell Biol 13, 79–85.

Orian, A., van Steensel, B., Delrow, J., Bussemaker, H.J., Li, L., Sawado, T., Williams, E., Loo, L.W., Cowley, S.M., Yost, C., et al. (2003). Genomic binding by the Drosophila Myc, Max, Mad/Mnt transcription factor network. Genes Dev 17, 1101–1114.

Orme, M.H., Alrubaie, S., Bradley, G.L., Walker, C.D., and Leevers, S.J. (2006). Input from Ras is required for maximal PI(3)K signalling in Drosophila. Nat Cell Biol 8, 1298–1302.

Pan, D., Dong, J., Zhang, Y., and Gao, X. (2004). Tuberous sclerosis complex: from Drosophila to human disease. Trends Cell Biol 14, 78–85.

Parker, J., and Struhl, G. (2015). Scaling the Drosophila Wing: TOR-Dependent Target Gene Access by the Hippo Pathway Transducer Yorkie. PLoS Biol 13, e1002274.

Perrimon, N., Pitsouli, C., and Shilo, B.Z. (2012). Signaling mechanisms controlling cell fate and embryonic patterning. Cold Spring Harb Perspect Biol 4, a005975.

Pesole, G., Gissi, C., De Chirico, A., and Saccone, C. (1999). Nucleotide substitution rate of mammalian mitochondrial genomes. J Mol Evol 48, 427–434.

Picton, H., Briggs, D., and Gosden, R. (1998). The molecular basis of oocyte growth and development. Mol Cell Endocrinol 145, 27–37.

Quinn, L.M., Dickins, R.A., Coombe, M., Hime, G.R., Bowtell, D.D., and Richardson, H. (2004). Drosophila Hfp negatively regulates dmyc and stg to inhibit cell proliferation. Development 131, 1411–1423.

Ren, Q., Zhang, F., and Xu, H. (2017). Proliferation Cycle Causes Age Dependent Mitochondrial Deficiencies and Contributes to the Aging of Stem Cells. Genes (Basel) 8.

Rios-Barrera, L.D., and Riesgo-Escovar, J.R. (2013). Regulating cell morphogenesis: the Drosophila Jun N-terminal kinase pathway. Genesis 51, 147–162.

Sarov, M., Barz, C., Jambor, H., Hein, M.Y., Schmied, C., Suchold, D., Stender, B., Janosch, S., K, J.V., Krishnan, R.T., et al. (2016). A genome-wide resource for the analysis of protein localisation in Drosophila. Elife 5, e12068.

Sieber, M.H., Thomsen, M.B., and Spradling, A.C. (2016). Electron Transport Chain Remodeling by GSK3 during Oogenesis Connects Nutrient State to Reproduction. Cell 164, 420–432.

Song, W., Ren, D., Li, W., Jiang, L., Cho, K.W., Huang, P., Fan, C., Song, Y., Liu, Y., and Rui, L. (2010). SH2B regulation of growth, metabolism, and longevity in both insects and mammals. Cell Metab 11, 427–437.

Stewart, J.B., Freyer, C., Elson, J.L., and Larsson, N.G. (2008). Purifying selection of mtDNA and its implications for understanding evolution and mitochondrial disease. Nat Rev Genet 9, 657–662.

Stewart, J.B., and Larsson, N.G. (2014). Keeping mtDNA in shape between generations. PLoS Genet 10, e1004670.

Stine, Z.E., Walton, Z.E., Altman, B.J., Hsieh, A.L., and Dang, C.V. (2015). MYC, Metabolism, and Cancer. Cancer Discov 5, 1024–1039.

Stump, C.S., Short, K.R., Bigelow, M.L., Schimke, J.M., and Nair, K.S. (2003). Effect of insulin on human skeletal muscle mitochondrial ATP production, protein synthesis, and mRNA transcripts. Proc Natl Acad Sci U S A 100, 7996–8001.

Templeman, N.M., and Murphy, C.T. (2018). Regulation of reproduction and longevity by nutrient-sensing pathways. J Cell Biol 217, 93–106.

Tiefenbock, S.K., Baltzer, C., Egli, N.A., and Frei, C. (2010). The Drosophila PGC-1 homologue Spargel coordinates mitochondrial activity to insulin signalling. EMBO J 29, 171–183.

Wallace, D.C. (2008). Mitochondria as chi. Genetics 179, 727–735.

Weston, C.R., and Davis, R.J. (2007). The JNK signal transduction pathway. Curr Opin Cell Biol 19, 142–149.

Zhang, Y., Chen, Y., Gucek, M., and Xu, H. (2016). The mitochondrial outer membrane protein MDI promotes local protein synthesis and mtDNA replication. EMBO J 35, 1045–1057.

Zhang, Y., Wang, Z.H., Liu, Y., Chen, Y., Sun, N., Gucek, M., Zhang, F., and Xu, H. (2019). PINK1 Inhibits Local Protein Synthesis to Limit Transmission of Deleterious Mitochondrial DNA Mutations. Mol Cell 73, 1127–1137 e1125.

## SUPPLEMENTAL REFERENCES

Bornstein, B., Zahavi, E.E., Gelley, S., Zoosman, M., Yaniv, S.P., Fuchs, O., Porat, Z., Perlson, E., and Schuldiner, O. (2015). Developmental Axon Pruning Requires Destabilization of Cell Adhesion by JNK Signaling. Neuron 88, 926–940.

Chen, D., and McKearin, D.M. (2003). A discrete transcriptional silencer in the bam gene determines asymmetric division of the Drosophila germline stem cell. Development 130, 1159–1170.

Edgar, R., Domrachev, M., and Lash, A.E. (2002). Gene Expression Omnibus: NCBI gene expression and hybridization array data repository. Nucleic Acids Res 30, 207–210.

Rhiner, C., Diaz, B., Portela, M., Poyatos, J.F., Fernandez-Ruiz, I., Lopez-Gay, J.M., Gerlitz, O., and Moreno, E. (2009). Persistent competition among stem cells and their daughters in the Drosophila ovary germline niche. Development 136, 995–1006.

Trcek, T., Lionnet, T., Shroff, H., and Lehmann, R. (2017). mRNA quantification using single-molecule FISH in Drosophila embryos. Nat Protoc 12, 1326–1348.

Wang, Z.H., Clark, C., and Geisbrecht, E.R. (2016). Drosophila clueless is involved in Parkin-dependent mitophagy by promoting VCP-mediated Marf degradation. Hum Mol Genet 25, 1946–1964.

Zhang, F., Qi, Y., Zhou, K., Zhang, G., Linask, K., and Xu, H. (2015). The cAMP phosphodiesterase Prune localizes to the mitochondrial matrix and promotes mtDNA replication by stabilizing TFAM. EMBO Rep 16, 520–527.

